# Optogenetic control of phosphate-responsive genes using single component fusion proteins in *Saccharomyces cerevisiae*

**DOI:** 10.1101/2024.08.02.605841

**Authors:** Matthew M. Cleere, Kevin H. Gardner

## Abstract

Blue light illumination can be detected by Light-Oxygen-Voltage (LOV) photosensing proteins and translated into a range of biochemical responses, facilitating the generation of novel optogenetic tools to control cellular function. Here, we develop new variants of our previously described VP-EL222 light-dependent transcription factor and apply them to study the phosphate-responsive signaling (*PHO*) pathway in the budding yeast *Saccharomyces cerevisiae*, exemplifying the utilities of these new tools. Focusing first on the VP-EL222 protein itself, we quantified the tunability of gene expression as a function of light intensity and duration, and demonstrated that this system can tolerate the addition of substantially larger effector domains without impacting function. We further demonstrated the utility of several EL222-driven transcriptional controllers in both plasmid and genomic settings, using the *PHO5* and *PHO84* promoters in their native chromosomal contexts as examples. These studies highlight the utility of light-controlled gene activation using EL222 tethered to either artificial transcription domains or yeast activator proteins (Pho4). Similarly, we demonstrate the ability to optogenetically repress gene expression with EL222 fused to the yeast Ume6 protein. We finally investigated the effects of moving EL222 recruitment sites to different locations within the *PHO5* and *PHO84* promoters, as well as determining how this artificial light-controlled regulation could be integrated with the native controls dependent on inorganic phosphate (P_i_) availability. Taken together, our work expands the applicability of these versatile optogenetic tools in the types of functionalities they can deliver and biological questions that can be probed.

## INTRODUCTION

Light is a ubiquitous environmental stimulus, harnessed by diverse organisms to control wide-ranging biological responses via photosensory proteins. Such proteins absorb light, triggering photochemical events at an internally-bound chromophore that are subsequently converted into conformational changes in the surrounding protein, culminating in altered downstream biological responses*^4^*.

In their simplest implementations, photosensors consist of two domains – a sensory domain and an effector – as exemplified by EL222, a 222-residue light-regulated transcription factor found in the marine bacterium *Erythrobacter litoralis HTCC2594^5, 6^*. The EL222 sensory component is a Light-Oxygen-Voltage (LOV) domain, a photosensory subset of the Per-ARNT-Sim (or PAS) superfamily of sensory proteins*^7, 8^*, which senses blue light via an internally-bound flavin chromophore. In addition, EL222 contains a single effector domain, a Helix-Turn-Helix (HTH) DNA binding domain. In the dark state, this protein adopts a “closed” monomeric structure, unable to bind DNA due to steric clashes preventing an obligate protein dimerization step. Upon blue light exposure, the flavin chromophore undergoes a photochemical reduction and concomitant adduct formation with a nearby cysteine residue, opening EL222 into a conformation exposing dimerizing surfaces in both the LOV and HTH domains, promoting DNA binding as a dimer*^5, 9^*.

Given its simplicity, reversible photoactivation, and rapid activation/deactivation kinetics, EL222 has been widely used as an optogenetic tool for transcriptional control in prokaryotes and eukaryotes*^10–15^*. In its original construction for eukaryotic applications, the first 13 residues of EL222 were replaced with a Nuclear Localization Sequence (NLS) and a VP16 Transcription Activation Domain (TAD), enabling “VP-EL222” to enter the nucleus and upon DNA binding, recruit RNA polymerase II to initiate transcription*^10^*. Parallel optimization of DNA binding sites by a combination of genomic mapping, SELEX (Systematic Evolution of Ligands by EXponential enrichment)*^16^*, and *in vitro* optimization led to the identification of a high affinity, non-native 20 bp binding sequence named “C120”. By arranging a five-fold tandem arrangement of this sequence (5xC120) in front of a minimal TATA promoter, we generated a potent and tunable system for increased gene expression under light activation*^10^*. Since this initial characterization, VP-EL222 has been used to control gene expression in multiple host systems including the budding yeast *Saccharomyces cerevisiae^10, 11, 17, 18^*, with the versatility provided by its precision tunability enabling investigations of diverse biological functions (e.g. characterization of fluorescent protein maturation) and new biotech applications via light-controlled expression of enzymes needed for microbial chemical production*^17, 19, 20^*.

Complementing this work with an artificial light-controlled promoter, *S. cerevisiae* has long been a workhorse for studies characterizing naturally-inducible promoters, as exemplified by the *PHO5* promoter*^21^*, a model for understanding how epigenetic control of gene transcription harnesses chromatin organization to differentially occlude cofactor binding sites*^22^*. *PHO5*, as well as the inorganic phosphate (P_i_) transmembrane transporter gene *PHO84*, is part of the larger phosphate-responsive signaling (PHO) pathway or *PHO* regulon that mediates the yeast phosphate stress response*^23^*. When phosphate is abundant, the promoter regions of *PHO5* and *PHO84* are nucleosome-bound*^1^*, blocking transcription factors such as Pho4 from accessing critical Upstream Activation Sequences (UAS) and the TATA box*^2, 3^*. As phosphate is depleted, dephosphorylated Pho4 (joined by Pho2) binds to a specific UAS, recruiting the SAGA (Spt-Ada-Gcn5 Acetyltransferase) complex to hyperacetylate histones in nearby nucleosomes*^24–26^*, causing them to unwind and reveal the previously-occluded binding sequences*^1, 2, 25, 27, 28^*. With these sites now accessible, transcription factors bind and drive expression of the *PHO5* and *PHO84* genes.

While conventional approaches have successfully characterized many aspects of *PHO5* and *PHO84* regulation, they have been unable to tackle certain questions due to fundamental limitations in existing tools. We believe that having access to reagents that can dynamically and precisely control gene expression are critical for fully manipulating transcriptional circuitry such as this, leading us to explore the versatility of VP-EL222 variants to better understand the nature of these chromatin driven promoters. In *S. cerevisiae*, some of the major complexes involved in chromatin remodeling and gene activation are SAGA (histone acetylation) and Rpd3S or Rpd3L (histone deacetylation). When SAGA is recruited to DNA by transcriptional activators such as Pho4, it binds and acetylates the histone tails of nearby nucleosomes through the Gcn5 histone acetyltransferase catalytic domain*^29–31^*, using acetyl-CoA for acetylation produced by the yeast acetyl-CoA synthetase ACS2 as a substrate*^32–34^*. For gene repression, the DNA-binding transcriptional regular Ume6 recruits Rpd3 histone deacetylation complexes to remove acetyl marks from nearby nucleosomes*^22, 35, 36^*.

In light of the roles of these diverse proteins and complexes in controlling gene expression, we pondered whether their light-dependent recruitment via EL222 could lead to a powerful new group of optogenetic transcriptional regulatory reagents. Here we report our first steps in developing such EL222 fusion constructs with a range of catalytic and non-catalytic components installed as N-terminal fusions to EL222’s LOV-HTH core. These effectors included catalytic domains involved in gene activation through chromatin regulation (Gcn5, Acs2, and Rpd3) as well as factors which recruit coactivator (Pho4) or corepressor (Ume6) complexes. We characterized the performance of these optogenetic regulators in *S. cerevisiae* in both plasmid-based and genomic settings, providing insights to the broader utility of these fusion proteins for probing two promoter regions of the yeast *PHO* regulon. Notably, EL222 variants containing Pho4 or Ume6 successfully activated and repressed target genes in predictable ways dependent on environmental factors (e.g. light, P_i_ concentration) and proximity to transcriptional start sites, demonstrating the utility of EL222-based approaches for up or downregulating targets in genomic contexts.

## RESULTS

### Tunability of EL222 in yeast using a single reporter plasmid

Our previously-published VP-EL222 optogenetic tool construct consists of NLS and VP16 sequences fused N-terminal to a truncated EL222 fragment (residues 14-222) which binds to the non-native, high affinity C120 sequence upon illumination (Figure S1a)*^10, 16^*. C120 can be arranged in a five-fold tandem repeat (5xC120) in front of a minimal TATA promoter for increased gene expression under light activation (Figure 1a). To investigate the temporal flexibility of VP-EL222 under various conditions, the Y422 yeast centromeric plasmid was developed for expression in *S. cerevisiae* under tryptophan deficient auxotrophic selection. This system utilizes VP-EL222 constitutively expressed by the *Ashbya gossypii* TEF fungal promoter, generating protein which binds elsewhere in the plasmid at the 5xC120 sequence upon blue light illumination, driving production of a yeast-optimized Yellow Fluorescent Protein (YFP) to quantitate EL222 activity (Figure 1b).

**Figure 1.**
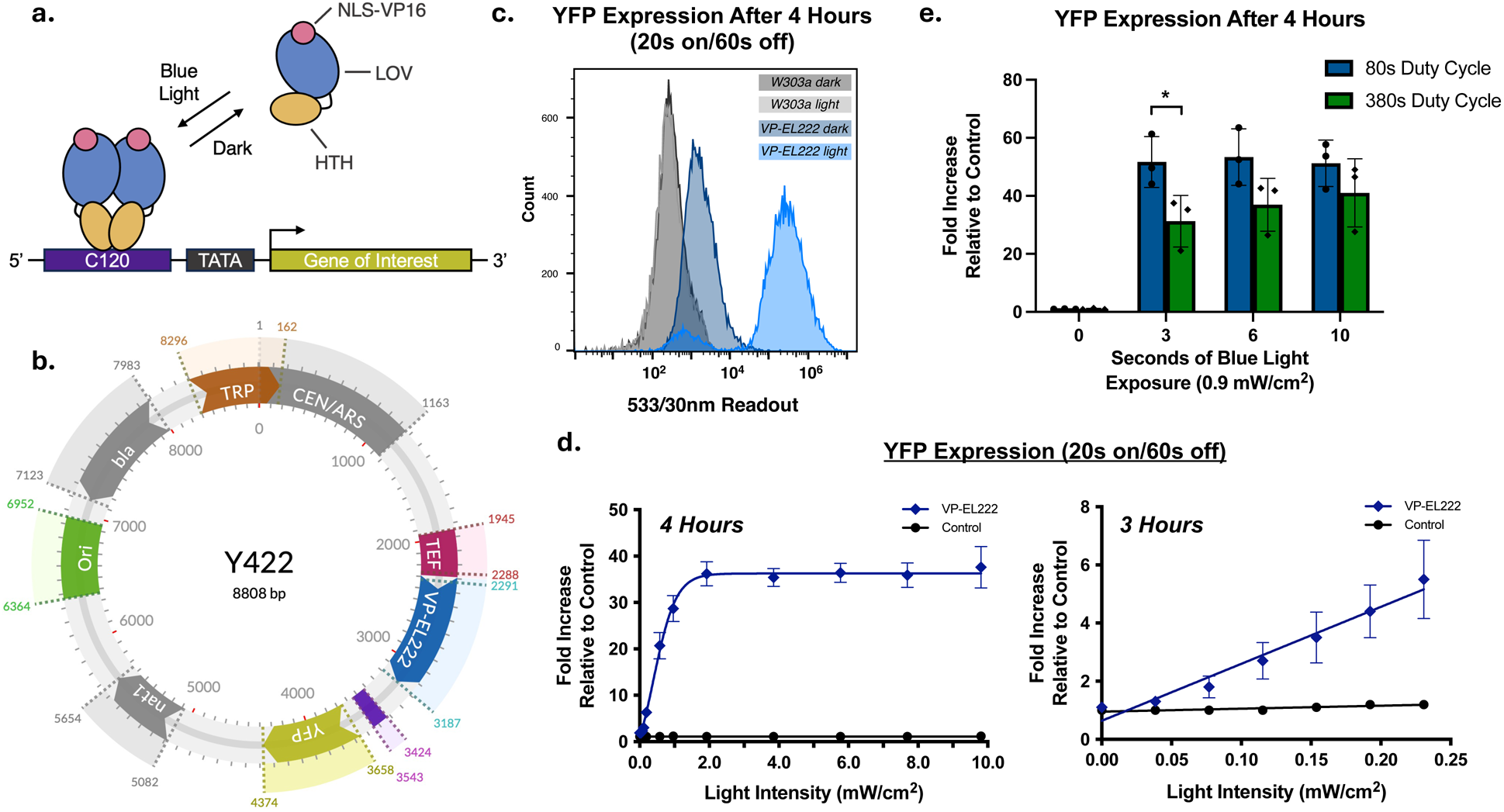
Plasmid-based expression of VP-EL222 driving Yellow Fluorescent Protein (YFP) production in *S. cerevisiae*. (a) Model of the blue-light activated transcription factor EL222 engineered for eukaryotic expression through addition of an N-terminal Nuclear Localization Sequence (NLS) and VP16 Transcription Activation Domain (TAD). In the dark, VP-EL222 is unable to bind DNA. Upon activation with blue light, VP-EL222 binds to the non-native C120 sequence (repeated five times), driving transcription of a downstream Gene of Interest (GOI). (b) Map of the yeast centromeric plasmid Y422 consisting of a constitutively active VP-EL222 (blue) driven by an *A. gossypii* TEF promoter (maroon), 5xC120 binding sequences (purple), YFP (yellow), origin of replication (green), tryptophan auxotrophic marker (brown), antibiotic resistance markers beta-lactamase and nourseothricin (gray), and the *S. cerevisiae* CEN4 centromere fused to the autonomously replicating sequence ARS1 (gray). (c) Flow cytometry histogram of W303a wildtype and Y422 transformed cells showing YFP activity in the dark and light after 4 hours (20 s on at 0.9 mW/cm^2^, 60 s off). (d) Tunable VP-EL222 activity reflected by YFP expression after blue light exposure (20 s on/60 s off). (e) VP-EL222 activity based on duty cycle variations (seconds of blue light exposure within a total cycle length outlined in the figure key). Data shown are mean ± SD. Significance was calculated using Two-Way ANOVA with Turkey’s multiple comparison test. \**p* ≤ 0.05.

To test the efficiency of this system, we used flow cytometry in the 533/30nm FITC channel to probe for YFP expression. As expected, *S. cerevisiae* W303a cells registered no change in fluorescence activity upon blue light activation, confirming that they are a suitable background for optogenetic testing. After transformation with the Y422 plasmid, we observed a minimal (∼5%) leakiness of VP-EL222 activity, consistent with our past experience*^10^* and many other optogenetic systems*^37^*. After 4 hours of pulsed blue light cycles (20 s on at 0.9 mW/cm^2^, 60 s off), we saw a >100-fold increase in VP-EL222 driven YFP activity (Figure 1c; gating approaches Figure S2). Similar results were seen with a plate reader-based readout, with YFP expression increased 30-fold (Figure 1d).

We quantified the intensity dependence of VP-EL222 driven gene expression in *S. cerevisiae* with a programable, in-house 96-well LED apparatus and titrating a gradient of light across each plate of cells (20 s on/60 s off). Using an 8-bit Arduino UNO microcontroller, we programmed each LED (with a *λ*_peak_ of 464.7 nm) from 0-255 intensity steps of 0.036 mW/cm^2^ each (Figure S3a-d). As light intensity strengthened, YFP expression increased until hitting 1.8 mW/cm^2^, where it registered a ∼36 fold increase relative to control after 4 hours. From there, YFP expression remained invariant to additional illumination, up to a maximum intensity of 9.2 mW/cm^2^. To develop a one-for-one relationship of light intensity to fold increase, we reduced both the total assay duration to 3 hours and focused on the lowest settings in the system. At the minimum intensities of ∼0.04-0.24 mW/cm^2^, a near linear pattern emerged (Figure 1d).

To understand the flexibility of time within each duty cycle, we explored both seconds of blue light exposure as well as length of intermittent darkness, keeping in mind the approximate 30 s time for the spontaneous reversion of EL222/chromophore structure and VP-EL222 driven gene expression we observed in mammalian cells*^5,10^*. Within our standard 80 s total cycle (20 s on/60 s off), shortening the illumination periods down to 10, 6, and 3 seconds (with corresponding lengthening of the dark periods) produced no discernable change. However, when we extended the total cycle duration from 80 s to 380 s, we began to see corresponding reduction of YFP expression, consistent with the dark deactivation time being substantially shorter than these longer dark periods. This became statistically significant at 3 s on/377 s off when compared to the 80 second cycles (Figure 1e). Further tests were performed to see the impact of light intensity when the total number of photons exposed to VP-EL222 remained the same. Short bursts of high intensity light outperformed extended periods of low light intensity, even in extended off cycles (Figure S4a-c). When we looked at the strength of single 60 s exposures, we observed a jump in YFP expression at the saturation point of 1.8 mW/cm^2^ out to 4 hours post-illumination (Figure S4d).

### Exploring the generality of EL222 fusions for light-dependent DNA recruitment

Compared to two-component systems, such as light-driven two-hybrid like associations of separated DNA binding and TAD activities, single protein systems like VP-EL222 are often easier to optimize and exhibit fewer technical drawbacks*^4, 38^*. However, a key variable of EL222- driven tools – their ability to tolerate fusions of molecular cargoes large than simple TADs – has not yet been rigorously tested to the best of our knowledge. To probe these constraints, we inserted a 26.7 kDa mCherry protein between the N-terminal NLS and TAD sequences and the LOV-HTH core in the Y422 background, letting us both examine the impact of this larger cargo on VP-EL222 function and enable the independent measurement of EL222 levels and activity via mCherry and YFP-selective fluorescence monitoring, respectively. Of note, we found that a substantial spacer between mCherry and the LOV-HTH core was essential for function, settling on a (GGGGS)_9_ linker in our final construct (Figure S1a). With this fusion protein in the reporter assays outlined in Figure 1 with a 20 s on/60 s off cycle for 4 hours at varying light intensities, we saw robust YFP production, indicating that photoreceptor function was unperturbed with the addition of this large protein. Additionally, our simultaneous observation of mCherry fluorescence provided strong evidence that exogenous domains can still properly fold and function while tethered to EL222.

To directly compare the performance of VP-mCherry-EL222 and VP-EL222 tools, we performed the optogenetic reporter assay at a light intensity of 3.6 mW/cm^2^ and saw comparable YFP expression to VP-EL222 with the 298 aa VP-EL222 and 595 aa VP-mCherry-EL222 demonstrating 43- and 44-fold increases using the plate reader, respectively (Figure 2a). Using widefield microscopy, we observed VP-mCherry-EL222 driven YFP production within the nuclei and cytosols of yeast cells. We also observed high levels of intracellular mCherry fluorescence, indicating proper functionality of this fluorescent protein (Figures 2b and S5). DAPI stained nuclei can be seen in the blue channel with their positions related to cellular fluorescence apparent when merged (Figure 2b).

**Figure 2.**
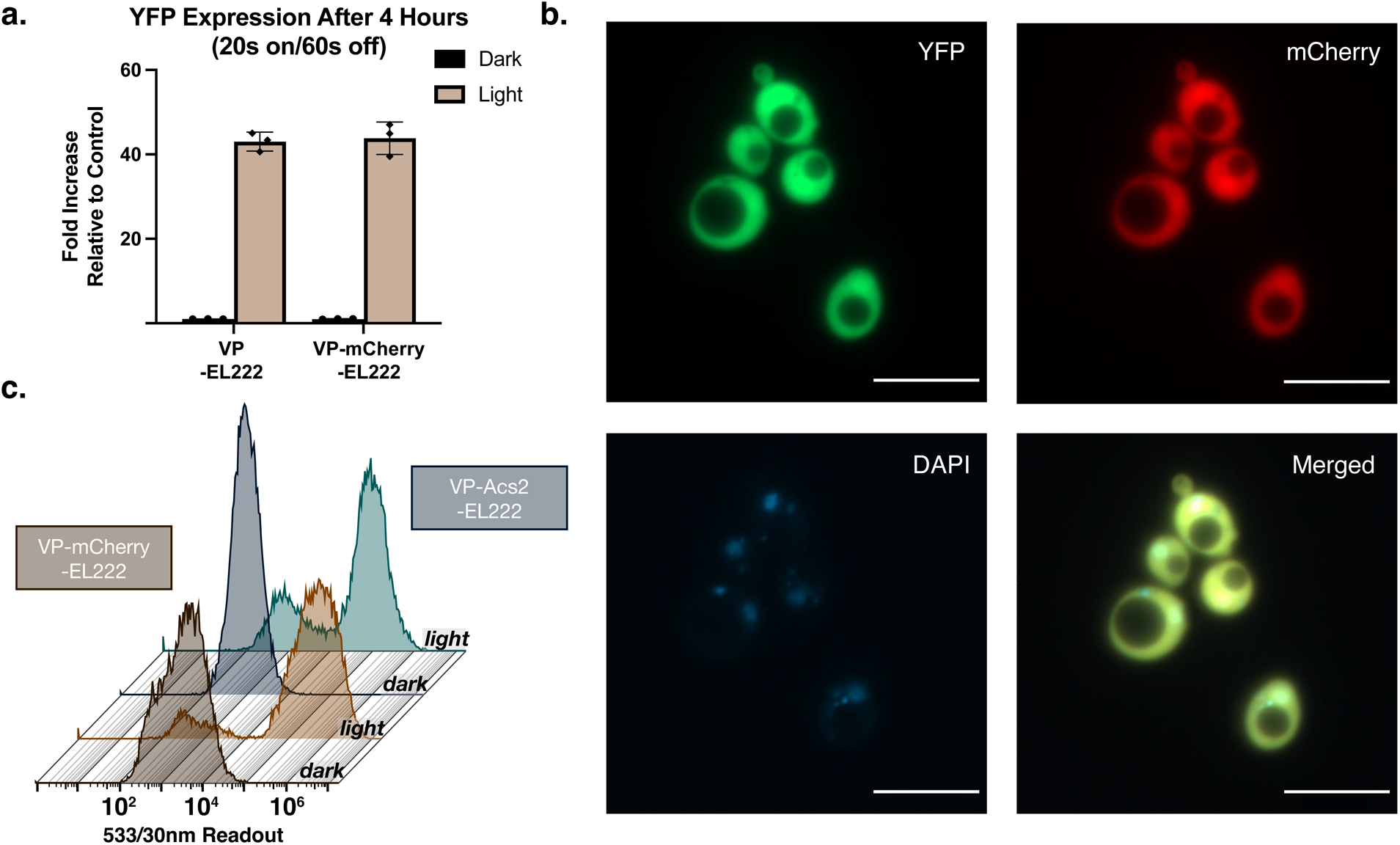
Relative activity of VP-EL222 fusion constructs with additional domains attached to the N-terminus of the LOV core through a flexible linker. (a) YFP activity of VP-EL222 and VP-mCherry-EL222 constructs after 4 hours (20 s on/60 s off) with a light intensity of 3.6 mW/cm^2^. (b) Wide-field microscope images of *Saccharomyces cerevisiae* cells displaying YFP production in the GFP channel (top left), mCherry activity (top right), DAPI staining of nuclei (bottom left), and merged layers (bottom right). *Scale bar = 10 µM*. (c) Flow cytometry histogram waterfall plots of VP-mCherry-EL222 and VP-Acs2-EL222 after 4 hours (20 s on/60 s off) with a light intensity of 0.9 mW/cm^2^.

We wanted to test the limits of our fusion constructs by replacing mCherry with effector domains we planned to utilize *in vivo*. The largest of these sequences was ACS2 (683 residues), substantially larger than mCherry and resulting in a 1,029 residue (111.2 kDa) VP- Acs2-EL222 complete fusion (Figure S1a). As with the smaller mCherry-containing construct, the inclusion of these large effector domains minimally impacted optogenetic gene expression, with VP-mCherry-EL222 producing a 117-fold increase in YFP expression versus a 75-fold increase for VP-Acs2-EL222 in a 4 hour fluorescence analyzer assay (Figure 2c; Figure S6b). Similar activity was also seen in the 784 residue VP-Gcn5-EL222 construct (Figure S6d). In contrast, we were unsuccessful in generating active VP-mCherry-Acs2-EL222 and VP-mCherry-Gcn5-EL222 constructs that exceeded the single domain-linker model, as no YFP expression or mCherry activity were observed with these systems (Figures S6c, e).

### Targeting yeast acid phosphatase genes using CRISPR and EL222 fusion constructs

With these optogenetic tools in hand, we proceeded to test them in genomic contexts starting with the *PHO5* gene, which encodes an enzyme responsible for phosphate hydrolysis important for cellular responses during periods of phosphate starvation*^39^*. Under high phosphate conditions (>300 µM), histones are tightly wrapped around the *PHO5* promoter region, obstructing both the TATA box and one of the two Upstream Activation Sequences (UAS)*^1, 40^*. This blocks the binding of the Pho4 transcription factor binds to the UAS, with an additional layer of repression caused by hyperphosphorylation-induced sequestration to the cytosol. As phosphate is depleted, Pho4 slowly dephosphorylates and becomes nuclear. Eventually, Pho4 can bind one of the two UAS (UASp1) (in association with Pho2) and recruit the SAGA complex to hyperacetylate surrounding histones*^24–26^*. These nucleosomes begin to unwind until the histones are ejected, exposing the second UAS site (UASp2) and TATA box. In this state, Pho4 binds UASp2 and induces transcription of *PHO5^1, 25, 41–43^*.

While *PHO5* responses to phosphate starvation are based on phosphatase production, a parallel response involves the upregulation of P_i_ transmembrane transporters encoded by the gene *PHO84^44^*. Like *PHO5*, *PHO84* is dependent on the transactivator Pho4 for activation, via a promoter that consists of 5 upstream activation sequences (UASpA-E) where UASpC/D act as high affinity sites with UASpB and UASpE occluded by upstream and downstream nucleosomes*^2^*. Upon phosphate depletion, these nucleosomes are ejected and *PHO84* transcription is induced (Figure 3a)*^1^*. However, it is important to note that *PHO5* and *PHO84* exhibit very different induction profiles with depleting levels of phosphate. As concentrations of inorganic phosphate (P_i_) are incrementally dropped from 250 µM to 0, *PHO84* undergoes a graded transition to activation where *PHO5* is more binary, staying nearly inactive until full depletion occurs*^42^*.

**Figure 3.**
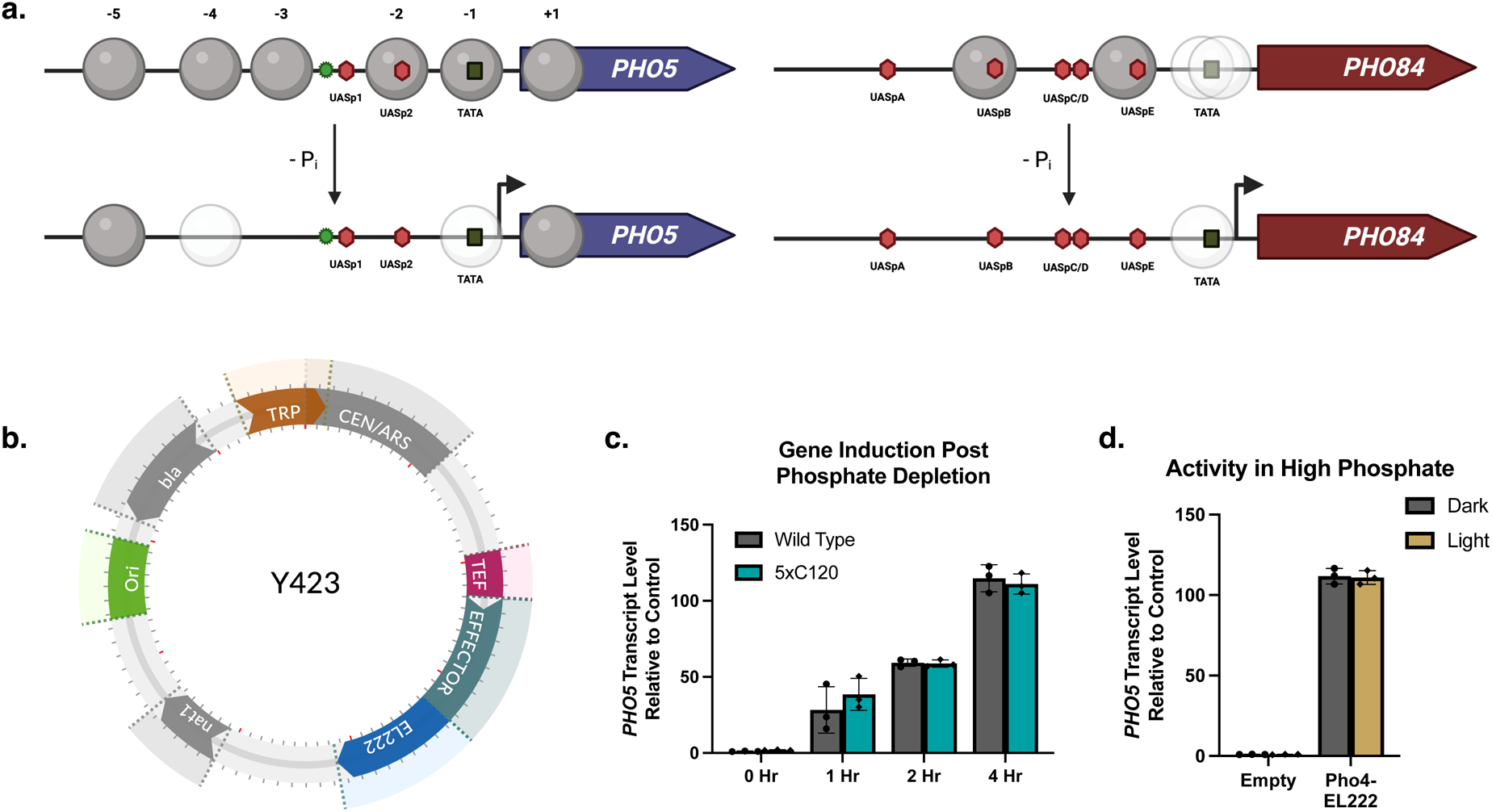
Induction of the yeast acid phosphatase genes *PHO5* and *PHO84* under native and non-native conditions. (a) In a high phosphate environment, some of the Upstream Activating Sequences (red hexagons) in the *PHO5* and *PHO84* promoters are blocked through steric occlusion by nucleosome forming histones. As inorganic phosphate (P_i_) is depleted, transcription factors such as Pho4 bind to the available UASp’s and initiate chromatin remodeling, leading to the eventual ejection of multiple histones and exposure of new UASp’s. More factors bind to these sequences eventually driving transcription of the *PHO5* and *PHO84* genes*^1–3^*. Dark spheres represent clearly positioned histones while faded spheres represent less organized nucleosomes. The green star indicates the initial insertion point of the 5xC120 binding sequence adjacent *PHO5*’s UASp1 site using CRISPR. (b) Generic map of the Y423 plasmids generated from the original Y422 plasmid. New effector domains are N-terminally fused to the LOV core through a flexible linker and the 5xC120- YFP reporter sequence has been removed. (c) Comparison of *PHO5* gene induction after phosphate depletion over 4 hours in both W303a wild type and 5xC120 promoter integrated cells. (d) Dark and light *PHO5* gene activity in high phosphate for the Pho4-EL222 construct and “Empty” NLS-EL222 control after 4 hours (20 s on at 15 mW/cm^2^, 60 s off).

To probe these genes with our newly-developed EL222 tools, we utilized CRISPR technology to stably integrate the 5xC120 binding sequence into the yeast genome (Figure S7). Then, we re-engineered the Y422 plasmid through our validations in Figure 2 into a plug-and-go construct named “Y423” for transient expression in the 5xC120 integrated background cells. For the Y423 constructs, the 5xC120 and YFP reporter sequences have been removed completely. Additionally, we replaced the VP16 TAD with a flexible linker engineered to facilitate fusing any effectors of our choice into the plasmid with standard restriction enzyme approaches (Figure 3b). Two of the catalytic domain constructs were tested in the Y422 plasmid background (20 s on at 0.9 mW/cm^2^, 60 s off) to confirm these effectors did not induce changes in gene expression through Pol II recruitment alone, and after 4 hours had produced no YFP activity in the light (Figures S8). With this approach, we produced constructs with five different effectors utilizing the multi-protein chromatin modifying complex recruiters *PHO4* and *UME6*, as well as the catalytic domains known to associate with them (*ACS2*, *GCN5*, and *RPD3*) (Figure S1b-c)*^45–48^*.

With these approaches in hand, we integrated the 5xC120 sequence near the initial Pho4 binding site in the *PHO5* promoter (green star in Figure 3a) and saw if it would alter the native expression of *PHO5* under depleted P_i_ conditions. Both wild type W303a and 5xC120 integrated cells were depleted of P_i_ and then observed for *PHO5* activation at various timepoints. mRNA levels were measured using quantitative PCR that were normalized to both ACT1 and ALG9 housekeeping genes. At 1 hour, 2 hour, and 4 hour timepoints, there was no significant difference in *PHO5* expression due to the insertion of the 5xC120 sequence. After 4 hours of gene induction, we observed an over 100-fold increase in *PHO5* transcript levels relative to control for both samples (Figure 3c).

We first deployed our catalytic domain constructs (Acs2, Gcn5, and Rpd3) under high P_i_ conditions (7.3 mM) targeting the *PHO5* promoter but were unsuccessful. No differences in ΔCt were observed between light and dark samples for all three constructs in relation to both the *ACT1* and *ALG9* housekeeping genes (Figure S9). These experiments were repeated in low phosphate media (50 µM P_i_) targeting either the *PHO5* or *PHO84* promoters, but the results were inconclusive or unsuccessful. When comparing the ΔCt of light vs dark, there was either no differences in relative expression when compared to the housekeeping genes, or the variability was inconsistent between *ACT1* and *ALG9* (Figures S10, S11). However, when we used the new transcription factor constructs Pho4-EL222 and Ume6-EL222, opposite phenotypes were observed. As seen in the mCherry constructs, the linker used to fuse proteins to the LOV core is long enough to allow proper folding and activity. With the addition of an NLS sequence to both Pho4-EL222 and Ume6-EL222, these proteins are quickly sequestered to the nucleus of each cell. In the case of Pho4-EL222, we observed *PHO5* transcript levels at 110- fold increase relative to the NLS-EL222 controls indicating perpetual *PHO5* activation regardless of illumination (Figure 3d). These levels were nearly identical to the phosphate depleted cells after 4 hours in Figure 3c. In high P_i_, Pho4 is heavily phosphorylated and kept cytosolic in yeast cells. Our data shows that both attachment to EL222 and the addition of an NLS sequence can overcome that natural repression, with enough room given by the linker for the attached Pho4 to bind to DNA on its own. By mutating a critical glutamic acid into leucine producing Pho4^E259L^-EL222, we successfully prevented this DNA binding in the dark. A similar issue arose with Ume6 also being active and forced nuclear when fused to EL222 but resulted in repeated lethality across all cells. After transformations in triplicate with Ume6-EL222, all plates were devoid of colonies. Ume6 is similar to the Zn2C6 GAL4 transcription factor, and so we used that protein as a guide to make a double mutation in critical cysteines responsible for DNA binding*^46, 49^*. Ume6^C770A/C787A^-EL222 was no longer able to bind to DNA on its own, providing successful transformation colonies (Figure S12).

### Activation and repression of the phosphate responsive gene *PHO5* by transcription factor, distance, and phosphate availability

First, we wanted to establish how our standard VP-EL222 was able to control gene function in a biologically simpler state. The chromatin landscape becomes much more accessible as phosphate is depleted, so we chose to lower the available P_i_ to a concentration of 50 µM. We used the Y421 yeast centromeric plasmid developed for expression in *S. cerevisiae,* with EL222 variants being constitutively expressed by an *A. gossypii* TEF fungal promoter (Figure S13). As a control for all *PHO5* transcript levels in Figure 4b-c, the dark state levels of VP-EL222 in high P_i_ at insertion point A was used. After 4 hours (20 s on at 15 mW/cm^2^, 60 s off), a well-defined pattern of expression for VP-EL222 along the *PHO5* promoter emerged. VP- EL222 had a maximum effect at the A site adjacent to UASp1 with *PHO5* transcript levels increasing by 67 fold after 4 hours. As we moved farther away from the UASp1 site, VP-EL222’s potency decreased, with *PHO5* transcript levels steadily dropping through sites B and C. At the farthest binding site D, the *PHO5* transcript levels remained unchanged with illumination (Figure 4b). When we raised the media [P_i_] to 7.3 mM, control over gene expression dramatically changed to a level of mixed success. At the initial sites A and B, VP-EL222 induced *PHO5* approximately 8- and 4-fold after 4 hours, respectively. As the distance increased to site C, we saw a spike up to 16-fold; while we saw the largest variability in data distribution here, all points were above the other samples. Site D again showed no change in *PHO5* transcript levels (Figure 4c). When comparing the dark state levels of *PHO5* activity, we observed at 4-5 fold decrease across all sites when the media [P_i_] was raised from 50 µM to 7.3 mM (Figure 4b-c).

**Figure 4.**
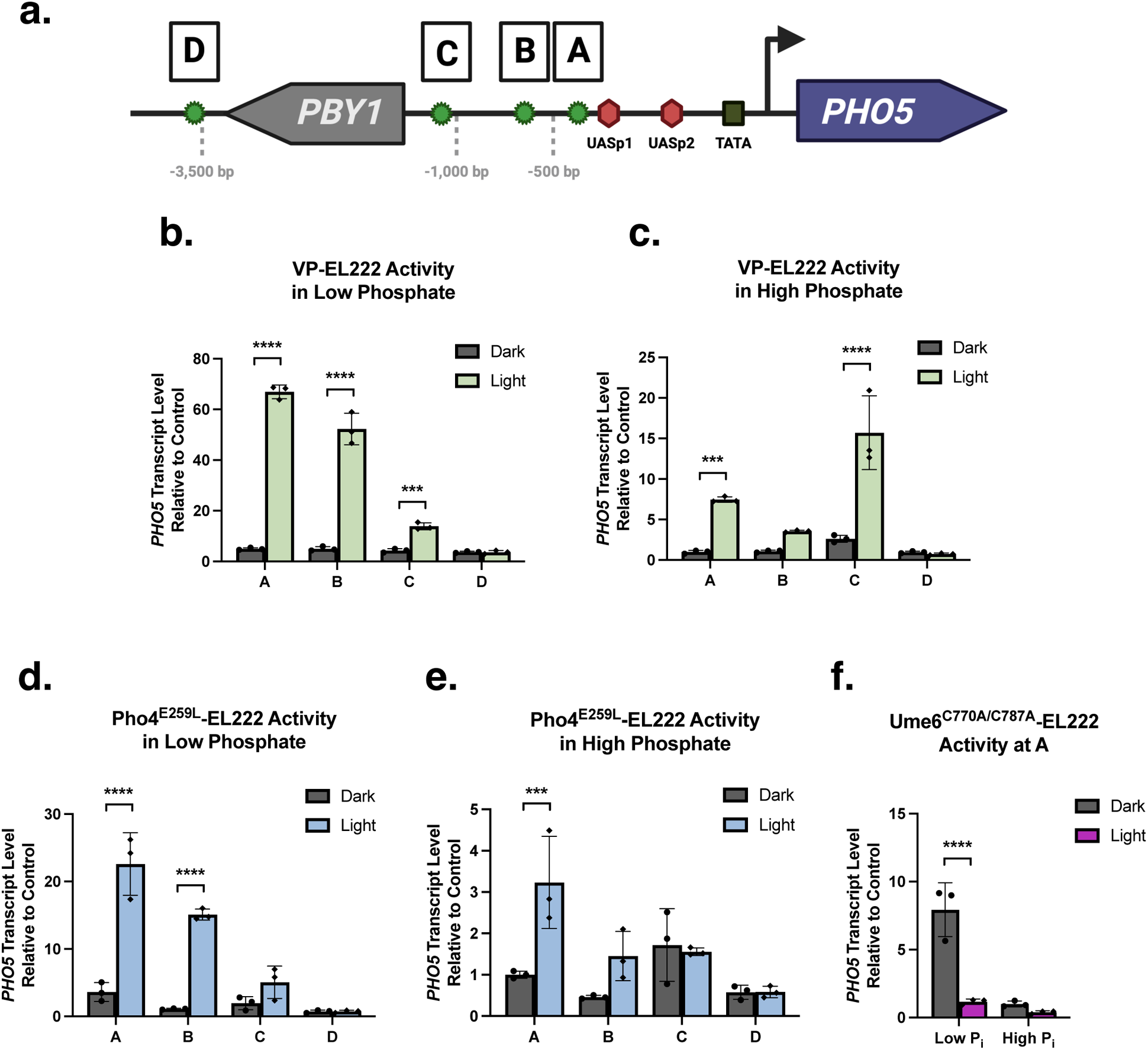
*PHO5* gene expression using various EL222 fusion constructs as related to both a function of distance and phosphate availability. (a) Representation of the *PHO5* promoter region with UAS sites (red hexagons) and the TATA box (black). Green stars indicate 5xC120 insertion sites along the *PHO5* promoter (A-D) with the farthest D site beyond the adjacent gene *PBY1*. (Distances from TSS: UASp1 - 361 bp, A - 414 bp, B - 560 bp, C - 1,029 bp, D - 3,510 bp). (b) VP-EL222 activity in low Pi (50 µM). (c) VP-EL222 activity in high Pi (7.3 mM). (d) Pho4^E259L^-EL222 activity in low Pi (50 µM). (e) Pho4^E259L^-EL222 activity in high Pi (7.3 mM). (f) Ume6^C770A/C787A^-EL222 activity in low and high Pi (50 µM and 7.3 mM, respectively). Data shown are mean ± SD. Significance was calculated using Two-Way ANOVA with Turkey’s multiple comparison test. \**p* ≤ 0.05, \*\**p* ≤ 0.01, \*\*\**p* ≤ 0.001, \*\*\*\**p* ≤ 0.0001.

While a highly potent TAD, VP16 is not at all biologically relevant to yeast, leading us to explore how EL222-based recruitment of a native activator like Pho4 could influence the *PHO5* expression. As a reference control for all *PHO5* transcript levels in Figure 4d-e, the dark state levels of Pho4^E259L^-EL222 in high P_i_ at insertion point A was used. Pho4^E259L^-EL222 was deployed in the same fashion as VP-EL222 and produced a defined comparable pattern under lower P_i_ conditions. At site A, a 23 fold increase in mRNA production is observed after 4 hours. This reduced in a stepwise fashion as we moved farther away along the promoter region through sites B and C. Once again, we saw no increase in *PHO5* transcript levels for site D (Figure 4d). When the phosphate levels are raised up to 7.3 mM P_i_, a similar pattern of variability emerged. Site A produced a 3 fold increase after 4 hours of blue light exposure. This dropped to negligible levels for sites B and C, with a wide distribution for dark state site C transcripts. Site D continued to show no effects here as well (Figure 4e). The broad distribution of sample points and less apparent patterns are similarly found for both VP-EL222 and Pho4^E259L^-EL222 in high P_i_. For dark state expression levels, we saw up to a 4 fold decrease across all sites when the media [P_i_] was raised from 50 µM to 7.3 mM (Figure 4d-e).

To complement the above optogenetic activation approaches, we wanted to see if similar strategies would allow us to repress gene expression levels as well. To do so, we utilized the Ume6 repressor, a key transcriptional regulator of early meiotic genes in yeast*^50^*. This subunit is a part of the greater Rpd3S/L histone deacetylase complex and does not normally bind in the *PHO5* promoter region*^51^*. Utilizing a double mutant in Ume6 predicted to disrupt inherent DNA binding in this protein and fusing it to EL222, we deployed Ume6^C770A/C787A^-EL222 into the A site only background cells containing the 5xC120 binding sequence. In both experiments, we used the dark state high P_i_ transcript levels as the control for *PHO5* mRNA expression. In low P_i_, the dark state basal activity of *PHO5* expression rose by an 8 fold increase. Through binding of Ume6^C770A/C787A^-EL222 at site A, we were able to repress expression of *PHO5* by nearly 8-fold back to the dark state high P_i_ transcript levels. But, under high P_i_ conditions, dark state expression levels were significantly reduced to the point Ume6^C770A/C787A^-EL222 was unable to depress them farther (Figure 4f).

### Activation and repression of the phosphate responsive gene *PHO84* by transcription factor, distance, and phosphate availability

To examine the general utility of these EL222-based tools, we investigated their performance in the similarly-related gene *PHO84*. While the promoter region of PHO84 is less understood than *PHO5*, *PHO84* contains five Pho4-binding UAS sites. We inserted 5xC120 sequences into two of these sites: AA (−230 bp from the TSS) and BB (−495 bp). Site AA occupies a space between UASpE and the TATA box (where nucleosomes are known to form) that should be accessible in a repressed state, while site BB was chosen because it sits between the UASpB (occluded by a histone), and the high affinity UASpC/D which are accessible in a repressed state (Figure 5a)*^2^*. For both low and high P_i_, we used the site AA dark state high P_i_ levels as the control reference for *PHO84* transcripts as it relates to each respective construct. First, VP-EL222 was used to target site AA under low P_i_ conditions. This produced a 9 fold increase in transcript levels when compared to the dark state. When we observed activity at site BB however, we saw no significant increase in levels. When we switched cells to high P_i_ (7.3 mM), we saw a 10 fold increase in transcript levels at site AA. In the light, both high and low P_i_ levels produced a wide distribution of datapoints with no significant difference between the two. As for site BB, there was no significant change in *PHO84* transcript production after 4 hours. When we looked at the dark state basal activity at 50 µM P_i_, site BB produced over a 6 fold increase, nearly 3 times as much as the site AA dark state basal activity. At the *PHO5* promoter, nearly all the dark state basal activity was the same, regardless of position. Here, because of the earlier activity and expression gradient of *PHO84* in response to phosphate levels, we observed a change in this activity. But, while site AA can still significantly boost expression of *PHO84* in the light, site BB was unable to provide an increase in transcription in both cases (Figure 5b).

**Figure 5.**
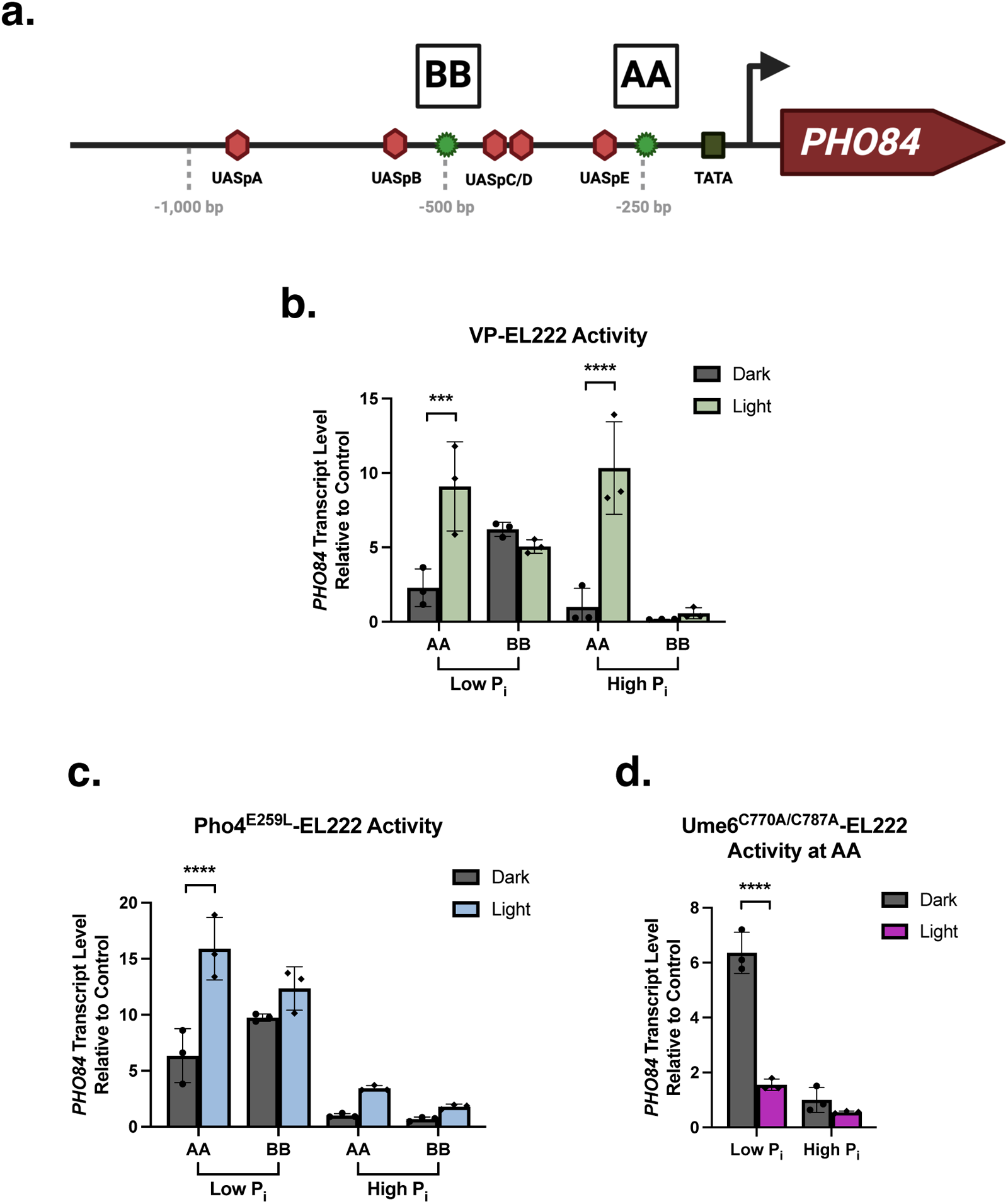
*PHO84* gene expression using various EL222 fusion constructs as related to both a function of distance and phosphate availability. Representation of the *PHO84* promoter region with UAS sites (red hexagons) and the TATA box (black). Green stars show 5xC120 insertion sites along the *PHO84* promoter (AA, BB). (Distances from TSS: AA - 230 bp, BB - 495 bp). (b) VP-EL222 activity in low (50 µM) and high (7.3 mM) P_i_. (c) Pho4^E259L^- EL222 activity in low (50 µM) and high (7.3 mM) P_i_. (d) Ume6^C770A/C787A^-EL222 activity in low (50 µM) and high (7.3 mM) P_i_. Data shown are mean ± SD. Significance was calculated using Two-Way ANOVA with Turkey’s multiple comparison test. \**p* ≤ 0.05, \*\**p* ≤ 0.01, \*\*\**p* ≤ 0.001, \*\*\*\**p* ≤ 0.0001.

Next, we moved on to the native transcription factor construct Pho4^E259L^-EL222. For site AA at low P_i_ levels, Pho4^E259L^-EL222 was able to produce a 16 fold increase of mRNA production, nearly double the value of VP-EL222 in Figure 5b. However, dark state activity was also higher registering a 6-fold increase when compared to the control. Site BB once again had the highest levels of dark state activity and showed no significant difference when exposed to the light. Under high P_i_ conditions, both sites AA and BB were unable to produce a significant increase in *PHO84* transcript levels after 4 hours of blue light exposure when compared to the dark start reference control (Figure 5c). The strength of the VP16 TAD was able to activate transcription at site AA under high P_i_ conditions when Pho4^E259L^-EL222 was unable to. Finally, we once again wanted to see if we could repress transcription of *PHO84* using Ume6^C770A/C787A^- EL222. We chose to target site AA with the dark state high P_i_ sample acting as the reference control. When P_i_ was depleted to 50 µM concentrations, the dark state levels of *PHO84* increased by over 6-fold. Under 4 hours of light activation, Ume6^C770A/C787A^-EL222 was able to successfully repress expression back to levels comparable to those seen in the dark state control. But, when the P_i_ concentrations were brought back up to 7.3 mM, gene expression was repressed and Ume6^C770A/C787A^-EL222 showed no impact (Figure 5d). While the variability of the *PHO84* promoter produced results from VP-EL222 and Pho4^E259L^-EL222 gene activation that were less apparent as the *PHO5* experiments, Ume6^C770A/C787A^-EL222 was shown to repress transcription in the same fashion throughout. At both *PHO5* and *PHO84*, our repression construct was able to depress transcript levels under active conditions all the way back to levels comparable to the high P_i_, repressed conditions.

## DISCUSSION

We better characterized VP-EL222 on a plasmid based system using *in vivo* analyses with multiple experimental devices for illumination and readout. A proof-of-concept for N-terminally tethered EL222 fusion proteins was a success after the addition of a substantial spacer between the extra domain and the LOV-HTH core. This led to the development of new transcriptional coactivator and corepressor EL222 fusion proteins native to the host system, *Saccharomyces cerevisiae*. We moved into genomic DNA targets through stable integration of the 5xC120 EL222 binding sequence using CRISPR, targeting the yeast phosphate stress response genes *PHO5* and *PHO84*. These chromatin driven promoter regions are accessible in a minimal phosphate environment but become increasingly more complex as phosphate becomes readily available. Initial probing with VP-EL222 was further extended by investigating these targets using non-VP effectors such as Pho4^E259L^-EL222 and Ume6^C770A/C787A^-EL222, leading to successful gene activation and repression.

We present tunable specifications for the blue-light activated transcription factor VP- EL222 in *Saccharomyces cerevisiae* using a single plasmid reporter system. A saturation point of ∼1.8 mW/cm^2^ was observed with a near one-to-one fold increase possible at the lowest end of the light intensity spectrum. We also confirm that pulse intensity is heavily weighted against number of photons over time as well as the off cycle length. This indicates that VP-EL222 is more resistant to change under cycling conditions and is much more tunable when focusing specifically on light intensities. A single 60 second exposure at maximum light intensity was able to produce over a 10-fold increase in YFP expression after 4 hours. This could be due to a cascading effect of active VP-EL222 at the 5xC120 promoter, the intensity of the VP domain as a TAD, and the maturation time of fluorescent proteins. Through N-terminal fusion of mCherry to the EL222 LOV-HTH core, we were able to confirm comparable activity to that of the VP-EL222 system and visualize proper functionality of the tethered protein. At 1,029 residues (111.2 kDa), the VP-Acs2-EL222 construct is one of the largest fusions of EL222 to be shown efficiently driving transcription. Tethering multiple domains together through a string of (GGGGS)_9_ linkers, however, was unsuccessful (Figure S6). These constructs mostly likely have become too complex for proper protein folding and activity.

These experiments became a proof-of-concept for EL222’s ability to translocate fused cargo to a specified region in the genome. We developed a plug-and-go plasmid system named “Y423” where effectors could be switched in and out with relative ease using standard molecular cloning techniques. Through CRISPR gene editing, the EL222 binding sequence 5xC120 was successfully integrated into various locations along the *PHO5* and *PHO84* promoters. Based on our initial assessments, these insertions had no effect on the native gene activation. We report here that the forced nuclear localization of Pho4 while fused to EL222 was enough to overcome the repressive phosphorylation under high phosphate conditions. This protein became active in the dark producing equal levels of *PHO5* transcripts as the wild type after 4 hours of depleted [P_i_] media. Interestingly, we were able to confirm previous observations of Ume6’s overexpression lethality through failed transformations*^52^*. Upon mutation of key cysteine residues involved in DNA binding, these cells became viable and were able to grow effectively.

The regulation and expression of *PHO* genes involved in the management of inorganic phosphate within the cell have always been stochastic in nature involving intrinsic noise*^1^*. We investigated their regulation optogenetically through multiple effector domains in fusions constructs with EL222 focusing on chromatin modifying complex recruiters and the catalytic domains known to associate with them. Unfortunately, our catalytic domain constructs consisting of Acs2, Gcn5, and Rpd3 were unable to alter gene activity at either promoter. We had hoped our sub-cellular localization of Acs2 to the chromatin would act in a similar manner to its mammalian homolog ACSS2*^53^* and assist in Acetyl-CoA production for histone acetylation but were unsuccessful. Some enzymes like Gcn5 are weak on their own and require the formation of subcomplexes for robust acetylation of nucleosomes*^31, 54^*. For our gene targets, these catalytic domains alone were insufficient to control gene function and mostly likely were unable to recruit other necessary components themselves. Multi-domain complexes are involved in chromatin remodeling, which is why when we deployed our transactivation domain constructs, we were successful in altering gene activity. We were able to show that under low P_i_ conditions, a clear and defined pattern of *PHO5* control emerged for both VP-EL222 and Pho4^E259L^-EL222. Sites closest to the UASp1 induced the highest levels of *PHO5* transcripts, with a descending gradient pattern forming as the 5xC120 sites were moved farther away. As the available phosphate was increased to 7.3 mM, the conditions become much more repressive. The complexity of the chromatin arrangement leads to variability in both accessibility and effectiveness at activating the system. Light induced VP-EL222 and Pho4^E259L^-EL222 formed little discernable patterns when activated across the promoter region, with a severely reduced capacity for transcription activation overall. In one case, VP-EL222 had the highest activation - 1,029 bp away from the TSS while Pho4^E259L^-EL222 was most effective adjacent the UASp1 site at −414 bp from the TSS. At a max distance of −3,510 bp located a full gene away, no effect can be seen on gene activity. We did observe a 4-5 fold increase of low P_i_ *PHO5* dark state activity across all sites when compared to the high P_i_ levels. This is probably due to some of the native nuclear Pho4 proteins becoming completely dephosphorylated and inducing low levels of activity*^40, 42^*. Hence, the chromatin state of the *PHO5* promoter is more accessible under low phosphate conditions and is why we see such a dramatic increase in coactivator driven transcript levels at sites A and B*^27^*. It is also worth noting, we do not know if the Pho4^E259L^ bound protein interacts with the nuclear Pho4 in low phosphate conditions. It is possible a knock-out of native Pho4 could reduce the *PHO5* expression seen here if this is the case.

The largest variability between VP-EL222 and Pho4^E259L^-EL222 was observed at the *PHO84* promoter, revealing the nuance between native and non-native effectors. Interestingly, VP-EL222 in the light produced the same levels of expression at site AA for both high and low P_i_. This is an alternative trend to the one observed up to this point for both VP-EL222 and Pho4^E259L^-EL222 at the *PHO5* promoter. Knowing that *PHO84* is more active at 50 µM P_i_, this indicates that VP-EL222’s activity at site AA is unaffected by chromatin structure. Conversely, Pho4^E259L^-EL222 followed the same trend in relation to phosphate availability as was seen in *PHO5*. As for the complete lack of site BB light-state activity, we still observe higher levels of dark state expression indicating the 5xC120 stable integration is most likely not the issue. Regardless of N-terminal effector used, this site is either unbound, negatively interacting with other DNA bound factors, or just is unable to drive transcription at this exact position regardless of P_i_ induced promoter arrangement. Pho4^E259L^-EL222 offers a native approach to gene activation and could be influenced by the other cofactors that interact with Pho4 protein within the cell. This is important when trying to understand complex chromatin systems like the *PHO* regulon with minimal invasiveness. While VP-EL222 was a stronger activator overall, this does not always make it the optimal choice as it lacks biological relevance to yeast. There have also been reports of VP16’s toxicity in certain organisms, requiring alternative options and highlighting the importance when choosing a transcription factor*^12^*.

Not only were we able to control gene activation using multiple coactivator fusion proteins, but we were also able to repress gene activity using the corepressor fusion construct Ume6^C770A/C787A^-EL222. While Pho4 is the traditional binding partner to the *PHO5* and *PHO84* promoters, Ume6 does not traditionally bind here. Ume6^C770A/C787A^-EL222 was able to repress gene activity under low P_i_ conditions back to the control levels seen in the high P_i_ dark levels. This up to 8-fold decrease indicates not only a return to an off state, but the potential at other promoters to repress at an even greater fold intensity. We attribute this to most likely occurring by Ume6^C770A/C787A^-EL222 recruiting the full histone deacetylase complex and actively repressing transcription through chromatin remodeling back to the off state. However, we cannot presently rule out this occurring instead through a passive mechanism, e.g. steric occlusion by the recruitment of certain Ume6-associated factors that simply block access to the transcription site.

In conclusion, we not only define a new set of parameters for novel engineered EL222 optogenetic tools, but also apply them strategically to investigate dynamic gene promoters. We present our plasmid based systems which can be manipulated using blue-light illumination in multiple experimental set-ups. With the stable insertion into any genome of the 5xC120 binding sequence, different effector-EL222 constructs can be transiently expressed with ease to target the same site under different environmental conditions. Here, we show the yeast *PHO5* and *PHO84* promoters are less definable using light induced gene activation under high P_i_, repressed conditions. This seems congruent with the stochastic nature of the chromatin landscape that controls the mechanisms of activity. These genes can be controlled by phosphate levels as well as distance, acting in a gradient, with activity based on strength of transcriptional activating effectors or repressors. *PHO84*’s gradient like response to phosphate levels and the complex nature of its promoter shows how multivariate gene expression can be, even in tightly control optogenetic experiments. Not only was our probing of the *PHO* genes a success, but this also continues to present a proof-of-concept for an EL222 plug-and-go system for a wide range of N-terminally fused effector domains.

## MATERIALS AND METHODS

### Yeast strains, media, and transformations

All *in vivo* experiments were performed using the budding yeast *Saccharomyces cerevisiae* ATCC 208352 strain W303-1a (gifted to us from Dr. Peter Lipke at Brooklyn College). Media used in this study was either YPD (1% yeast extract, 2% peptone, 2% glucose) supplemented with adenine (4 g/L), tryptophan (8 g/L), and uracil (2 g/L) or standard synthetic media (complete, tryptophan, or uracil deficient). For the phosphate mediated experiments, yeast nitrogen base without amino acids and phosphate (Formedium CYN08**) was used to create standard synthetic media (complete or tryptophan deficient) which could then be supplemented with the experimental concentrations of phosphate (0 µM, 50 µM, and 7.3 mM).

For transformations, *S. cerevisiae* W303a cells were plated on YPD agar (2% agar) and grown at 30 °C. 30 mL cultures of YPD broth were inoculated using a single colony from each plate and grown overnight at 200 rpm/30 °C. Cultures were diluted in the morning to an OD_600_ of 0.2 in 50m mL YPD broth and left to grow at 200 rpm/30 °C until an OD_600_ of 0.4-0.5 was reached (∼2-3 hours). Cells were then harvested by centrifugation (4,000 rpm/ 5 min), washed with 20 mL of ice-cold sterilized ddH_2_O, and centrifuged again (4,000 rpm/ 5 min). Cell pellets were resuspended in a 20 mL solution of 0.1 M Lithium Acetate, 0.8 M Sorbitol, and 1xTE buffer for 30 min at 30 °C followed by the addition of 0.2 mL of 1 M DTT for 15 min at 30 °C. Cells were centrifuged (4,000 rpm/5 min), washed twice with 20 mL of ice-cold 1M sorbitol, and then resuspended in 0.2 mL ice-cold 1 M sorbitol.

For each yeast transformation by electroporation, 45 μL of cell suspension was combined with 5 μL of plasmid (400 ng/μL;∼10kb) in a microfuge tube and left to incubate on ice for 15 min For CRISPR gene editing, 4 μL of plasmid (400 ng/μL;∼10kb) and 6 μL of DHR cassette was used with 40 μL of cell suspension was used instead. Cell suspensions were transferred to 0.2 cm electroporation cuvettes and electroporated using a BioRad Gene Pulser xCell on its pre-set *S. cerevisiae* protocol (1.5 kV, 200 mA, 25 μF). Post electroporation, the cells were immediately resuspended in 1 mL of 1 M sorbitol and left to recover at 30 °C for 1 hour. Cells were centrifuged (4,000 rpm/5 min), resuspended in 200 μL 1 M sorbitol, and spread on tryptophan or uracil deficient synthetic drop out agar for selection based on the plasmid used.

### Plasmids, assembly of fusion constructs, and CRISPR genome editing

Yeast centromere plasmids Y421 and Y422 (provided by Farren Isaacs, Yale University) were utilized with tryptophan auxotrophic markers (Figures S13 and 1b, respectively). Y421 and Y422 both contain a constitutively active TEF fungal promoter that drives EL222 production, but Y422 contains an additional 5 tandem repeat of EL222’s (UniProt ID: Q2NB98) DNA binding sequence C120 (5XC120) upstream of the Yellow Fluorescent Protein (YFP) sequence*^10^*. Y422 was used to make all the fusion constructs through primer design and PCR mediated sequencing editing. Additions to the plasmid were made through targeted PCR of the backbone plasmid and desired insert followed by restriction enzyme digestion and ligation using Quick Ligase (NEB Catalog #M2200). Plasmid backbones were extracted using a Monarch Gel Extraction kit (NEB Catalog #T1020). Effector domain sequences (ACS2, GCN5, RPD3, PHO4, and UME6) were taken straight from the host yeast cells genomic DNA by PCR for ligation in Y422. Targeted residue mutations within effector domains (Figure S1b-c) were produced via site-directed mutagenesis*^55, 56^* followed by DpnI digest and ligation with Quick Ligase (NEB Catalog #M2200). Alternately, a standard mCherry sequence was used in the double fluorescence constructs (Table S1; Supplemental Sequence S1).

All linkers, scaffolds, and CRIPSR HDR cassettes were produced either through gene fragment synthesis ordered from Twist Biosciences, or through PCR, RE digest, and ligation into the plasmid pHisGB1. Desired sequences where then amplified from these templates using PCR. Subtractions from these plasmids was performed using PCR primers designed with restriction enzyme sites for partial plasmid sequence amplification followed by restriction enzyme digest (including DpnI) and ligation. Final constructs were produced through the removal of Y422’s 5xC120-YFP sequences creating a set of plasmids named Y423 (Figure 3b). Y421 had its VP16 sequence removed to create the NLS-EL222 “Empty” construct.

For CRISPR gene editing, the “plug-and-go” plasmid pML104 was ordered from Addgene (a gift from John Wyrick [Washington State University] Addgene plasmid #67638; http://n2t.net/addgene:67638; RRID:Addgene_67638) containing the Cas9 protein, guide RNA expression cassette with BclI-SwaI cloning sites for guide sequence cloning, and a URA3 uracil marker for yeast transformation*^57^*. The 20 bp guide RNA sequences were designed using CHOPCHOP*^58–60^*, placed in oligonucleotide sequences, and ordered from IDT. Oligonucleotides were annealed in a thermocycler to themselves and used as inserts into the pML104 plasmid using Quick Ligase (Table S1). For stable integration of the 5XC120 EL222 binding sequence into the *PHO5* and *PHO84* promoter regions, DHR cassettes were designed with the desired sequence sandwiched between homology arms (∼150 - 300 bp each) identical to the genomic DNA flanking the insertion point (Table S1). pML104 and the corresponding DHR cassette were co-transformed into yeast cells where the insert was integrated into the genome via double homologous recombination (Figure S7). All plasmid described here were proliferated in *Escherichia coli* DH5α cells and subsequently purified with a Promega Wizard Genomic DNA Purification Kit (Catalog #A1120). All plasmid alterations, ordered inserts, and CRISPR genomic editing was confirmed through Sanger sequencing (GENEWIZ).

### Genomic DNA and RNA extractions

For genomic DNA extractions, yeast cells were grown overnight (200 rpm/30 °C) to a required density of OD_600_ 1.0-2.0. Cells were pelleted in a 2 mL Eppendorf tube, the supernatant discarded, and the cell pelleted resuspended in 1.0 mL of DNA extraction buffer (50 mm Tris HCl, 10 mM EDTA, 50 mM NaCl, 1% [w/v] SDS, pH 7.5). “250 µL” (by volume) of sterile glass beads 425-600 μm (Sigma G8772) was added and vortex for 30 seconds to break up the yeast cells followed by incubation at 65 °C for 30 minutes. Samples were centrifuged (13,000 rpm/10 min) and 600 µL of supernatant was removed and placed in a new tube. 600 µL of phenol:chloroform:isoamyl alcohol (25:24:1 v/v/v) was added, inverted several times to mix, and centrifuged (13,000 rpm/10 min). The upper aqueous phase was removed, mixed with 0.7 volumes of cold isopropanol in a new tube, and incubated at - 20 °C for 1 hour. Samples were centrifuged (13,000 rpm/10 min), supernatant discarded, cell pellet washed with 700 µL of 70% ethanol and centrifuged again (13,000 rpm/10 min). The 70% ethanol was removed, and the cell pellet was allowed to dry in the fume hood before being resuspended in sterilized ddH_2_O.

For RNA extractions, cells were flash frozen until every sample was ready for extraction. Cells were thawed on ice and the RNA was extracted using a RiboPure Yeast RNA purification kit (Invitrogen; Catalog #AM1926) as per the manufacturer’s instructions. RNA samples were then flash frozen in liquid N_2_ for later use in cDNA production and qPCR analysis.

### Blue light illumination of samples

All *S. cerevisiae* strains were grown in 20 mL liquid cultures inoculated with single colonies and grown overnight in total darkness (200 rpm/30 °C). Cultures were diluted in the morning to an OD_600_ of 0.2 and left to grow at 200 rpm/30 °C until an OD_600_ of 0.3-0.4 was reached (∼2 hours). Cell cultures were then distributed in an identical pattern into either two Corning Black with Clear Flat Bottom 96-well plates at 100 µL per well or Clear 24-well plates at 800 µL per well. One plate was covered with aluminum foil and set aside as a dark control. For the 96-well plates only, each experiment was placed on a custom built 96-well LED system (courtesy of M. Bikson, CCNY Department of Biomedical Engineering) in the dark and covered with a red-coated piece of board. 96 individual WS2812B LEDs fed into fiberoptic wires deliver independently controllable light conditions controlled by an Arduino UNO microcontroller. Blue light is emitted from these LEDs at a peak wavelength of 464.7 nm (read by an OceanOptics spectrophotometer) with a light intensity of 0.04-9.22 mW/cm^2^ (Figure S3). The light plate was exposed to repeating cycles of blue light (20 s on/60 s off) for up to 4 hours. For the phosphate based experiments, a blue LED panel (465 nm, 13.8 Watts; 2501BU - LED Wholesalers) was suspended ∼6 inches from the top of the 24-well plates with a light intensity of 15 mW/cm^2^ and set to repeating cycles of blue light (20 s on/60 s off) using an electronic intervalometer (Model 451, GraLab) (Figure S14).

### Fluorescence analysis by plate reader, flow cytometry, and microscopy

For whole well reads, a SpectraMax i3 multimode microplate reader (Molecular Devices) was used to analyze fluorescence intensity. Cells were first resuspended using a multi-channel pipette before loading into the plate reader and the data was analyzed using GraphPad Prism software. For flow cytometry, samples were analyzed using a BD Accuri C6 Sampler Plus flow cytometer, light plate first, followed by unwrapping of the dark plate. All wells were resuspended using a multi-channel pipette at a maximum of 6 wells per read to prevent variations in cell sedimentation. 20,000 events were collected per well at a threshold of 50,000 under slow flow.

Standard gating techniques were used to select single-cell yeast populations and applied to subsequent FITC channel (533/30 filter) histograms used to detect the presence of YFP (Figure S2). Data was then processed using FlowJo software*^61^*. mCherry fluorescence was observed using either a wide-field Axio Observer.Z1/7 Inverted Microscope (Zeiss) with a Plan-Apochromat 63x/1.40 numerical aperture (NA) oil objective or a Marianas Spinning Disk confocal microscope (Intelligent Imagine Innovations) consisting of a spinning disk confocal head (CSU-X1, Yokagawa) on a Zeiss Axio Observer inverted microscope equipped with x100/1.46 numerical aperture (NA) Plan-Apochromat (oil immersion). Frames were captured on an Axiocam 506 mono (Zeiss) using Zen software (Zeiss) for the wide-field and a Prime sCMOS camera (Photometrics) controlled by SlideBook 6 (Intelligent Imagine Innovations) for the spinning disk confocal and further processed using ImageJ*^62, 63^*. DAPI staining was performed using a final concentration of 10 µg/mL added to mid log-phase cells (OD_600_ of ∼0.5) and let sit for 10 minutes. Cells were then harvested by centrifugation (4,000 rpm/5 min), washed with 1xPBS, centrifuged again (4,000 rpm/5 min), and resuspended in 1xPBS before imaging.

### EL222 fusion protein experiments at the *PHO5* and *PHO84* promoters with quantitative PCR

All 5xC120-*PHO5* and 5xC120-*PHO84* integrated *S. cerevisiae* strains containing their respective fusion constructs were grown in 20 mL liquid cultures inoculated with single colonies and grown overnight in total darkness (200 rpm/30 °C). Cultures were diluted in the morning to an OD_600_ of 0.2 and left to grow at 200 rpm/30 °C until an OD_600_ of 0.3-0.4 was reached (∼2 hours). Cell cultures were then distributed in an identical pattern into two Corning Black with Clear Flat Bottom 96-well plates at 100 µL per well or Clear 24-well plates at 800 µL per well. One plate was covered with aluminum foil and set aside as a dark control. The other plate was illuminated as outlined previously. Samples were collected and flash frozen in liquid N_2_ for later use. Samples were then thawed, and the RNA was extracted from each sample as outlined previously.

RNA for each sample was diluted to a total concentration of 500 ng in 16 µL with 4 µL of qScript Ultra SuperMix (Quantabio #95217) added for a total of 20 µL. Samples were run in a thermocycler for a single cycle (2 min/25 °C, 10 min/55 °C, 1 min/95 °C) to create 500 ng of cDNA. These samples were diluted down to a concentration of 1 ng/µL and distributed into a 384-well plate in triplicate (5 µL per well). Genes are targeted using primer pairs to produce sequence lengths of ∼100-300 bp (*PHO5* or *PHO84* and two housekeeping gene controls *ACT1* and *ALG9*) (Table S1)*^64–66^*. Both forward and reverse primers were mixed in the same tube to a stock concertation of 10 µM. 7 µL of master mix (6 µL PerfeCTa SYBR Green FastMix [Quantabio #95071], 0.5 µL primer stock, 0.5 µL dd H_2_O) was added to each sample well for a total volume of 12 µL per well. The plate was then sealed and analyzed using a QuantStudio 7 Flex Real-time qPCR system. For experiments performed in biological triplicates, data was analyzed using the 2^-ΔΔCT^ method with the geometric mean of both *ACT1* and *ALG9* housekeeping gene controls. For the technical replicates of n = 1, data was analyzed using the ΔCT of each housekeeping gene control.

## ASSOCIATED CONTENT

### Supporting Information

Supplemental Figures S1-S14, Table S1, and Sequence S1 are provided as a compilation of in-depth resources to assist in any technical questions based on the main publication. This includes outlines of all the constructs used in this study along with a reference sequence, supporting flow cytometry/qPCR data, and technical specifications for major points provided in the text.

## AUTHOR INFORMATION

### Author Contributions

M.M.C.: conceptualization, investigation, formal analysis, writing; K.H.G.: conceptualization, supervision, editing, funding.

### Notes

The authors declare no competing financial interest.

## Supporting information

Supplemental Sequence File S1

## ACKNOWLEDGEMENTS

We thank Farren Isaacs (Yale University, Yale School of Medicine) for gifting the Y421 and Y422 plasmids and Peter Lipke (Brooklyn College, School of Natural and Behavioral Sciences) for supplying the W303a *S. cerevisiae* strains used in this study. We thank Marom Bikson and Robin Azzam (CCNY, Department of Biomedical Engineering) for the custom 96-well LED machine. We thank Alfredo Vidal Ceballos and Rachel Fisher (CUNY Advanced Science Research Center) for assistance with the wide-field and confocal microscopy. We would also like to thank Jia Liu, Catherine Leckie, and Sami Sauma for their assistance with quantitative PCR. Graphical abstract and Figures 3a, 4a, 5a, and S7 were all created with BioRender.com. Plasmid maps found in Figures 1b, 3b, and S13 were all created with AngularPlasmid.

