## Supplemental Information for "Optogenetic control of phosphate-responsive genes using single component fusion proteins in *Saccharomyces cerevisiae*"

Contents:

Figures S1-S14
Table S1
Supporting References

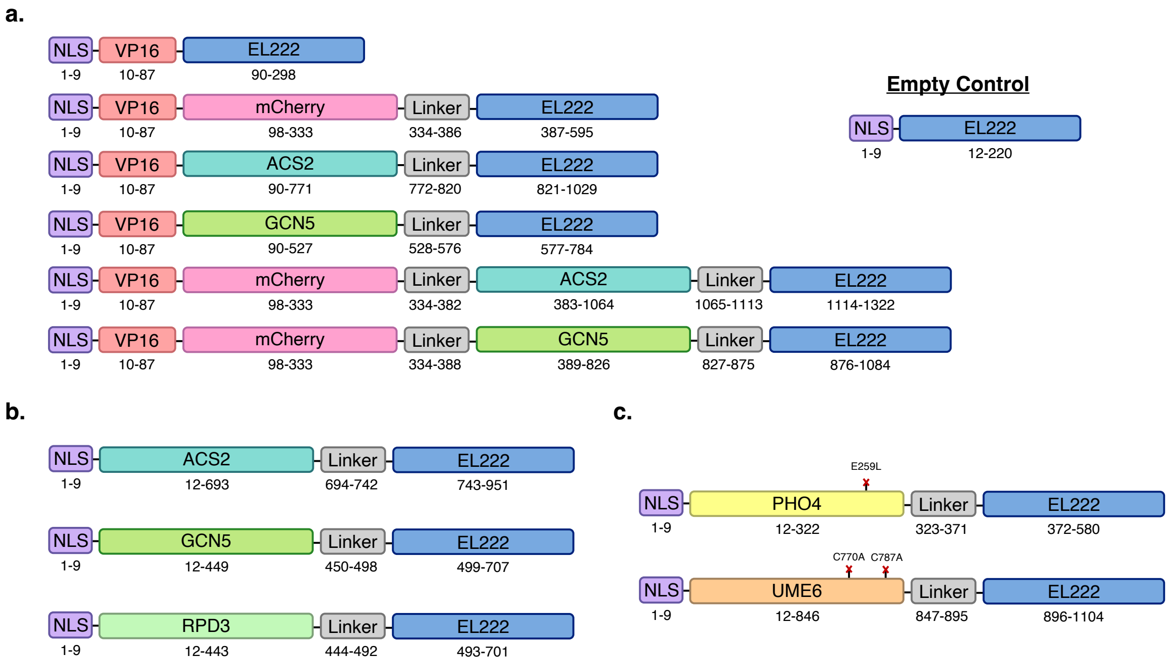


**Figure S1.** Major constructs used in this study. (a) VP16 based constructs with the original "EL222" consisting of an NLS, VP16, and EL14-222 sequence at the top. $(GGGGS3)_9$ flexible "Linkers" are used to fuse various domains. "Empty" control construct consists of the NLS and EL222 sequences only. (b) Fusion constructs consisting of domains known to be involved in histone acetylation/de-acetylation. The VP16 TAD has been removed to prevent direct transcription initiation through Pol II recruitment. (c) Fusion constructs consisting of native *S. cerevisiae* transcription factors with the VP16 TAD removed. $PHO4^{E259L}$ and $UME6^{C770A/C787A}$ constructs were produced in addition to the wild type sequences, preventing each domain from directly binding DNA on their own (marked by red x's)[1, 2].

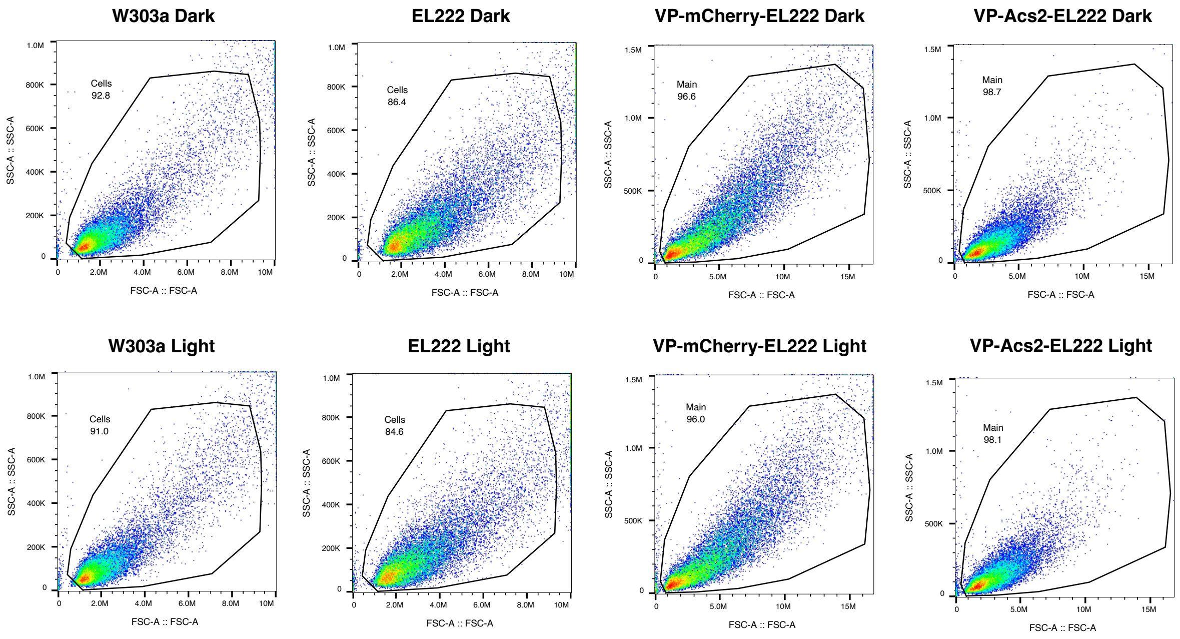


**Figure S2.** Flow cytometry density scatter plots showing the gating strategy for Figs. 1 and 2. Side scatter over forward scatter data was gated to isolate normal yeast cells from background noise. The same gating approach was taken for Supplemental Figures S6 and S8.

a.

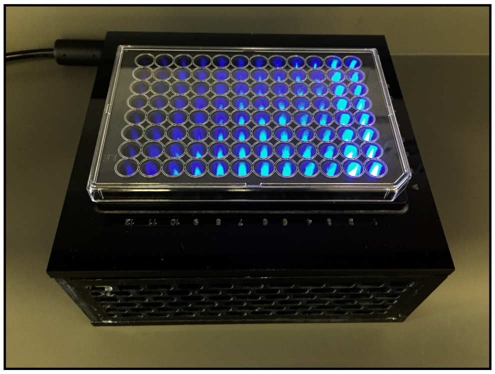

b.

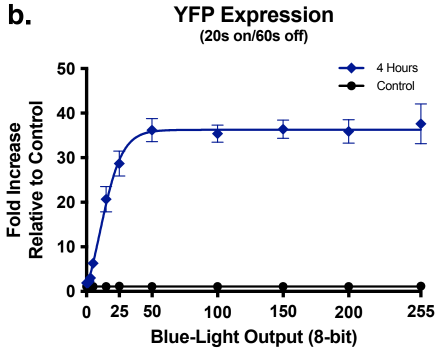


c.

| LED Color | $\lambda_{peak}$ (nm) | FWHM (nm) |
|---|---|---|
| Red | 632.9 | 17.4 |
| Green | 521.5 | 39.8 |
| Blue | 464.7 | 25.3 |

d.

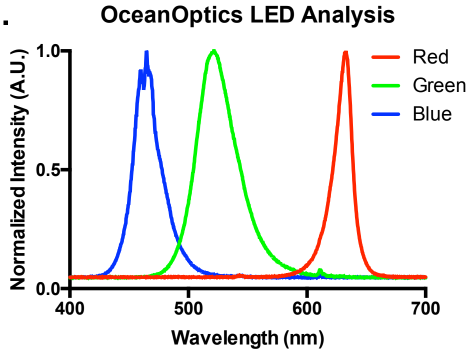


**Figure S3.** In-house built LED machine for optogenetic experiments. (a) 96 individual LED's emit light that is fed into the bottom of a 96 well plate using fiber optic cables. The LED panel is controlled by an Arduino UNO microcontroller with independent regulation over multiple wells for titration experiments. (b) YFP expression of the Y422 plasmid in *S. cerevisiae* in relation to the 8-bit settings for blue-light output. Each integer produces a light intensity of 0.036 mW/cm$^2$. (c) RGB $\lambda_{peak}$ (nm) and FWHM (nm) LED values. (d) OceanOptics LED analysis RGB histogram (as it relates to c).

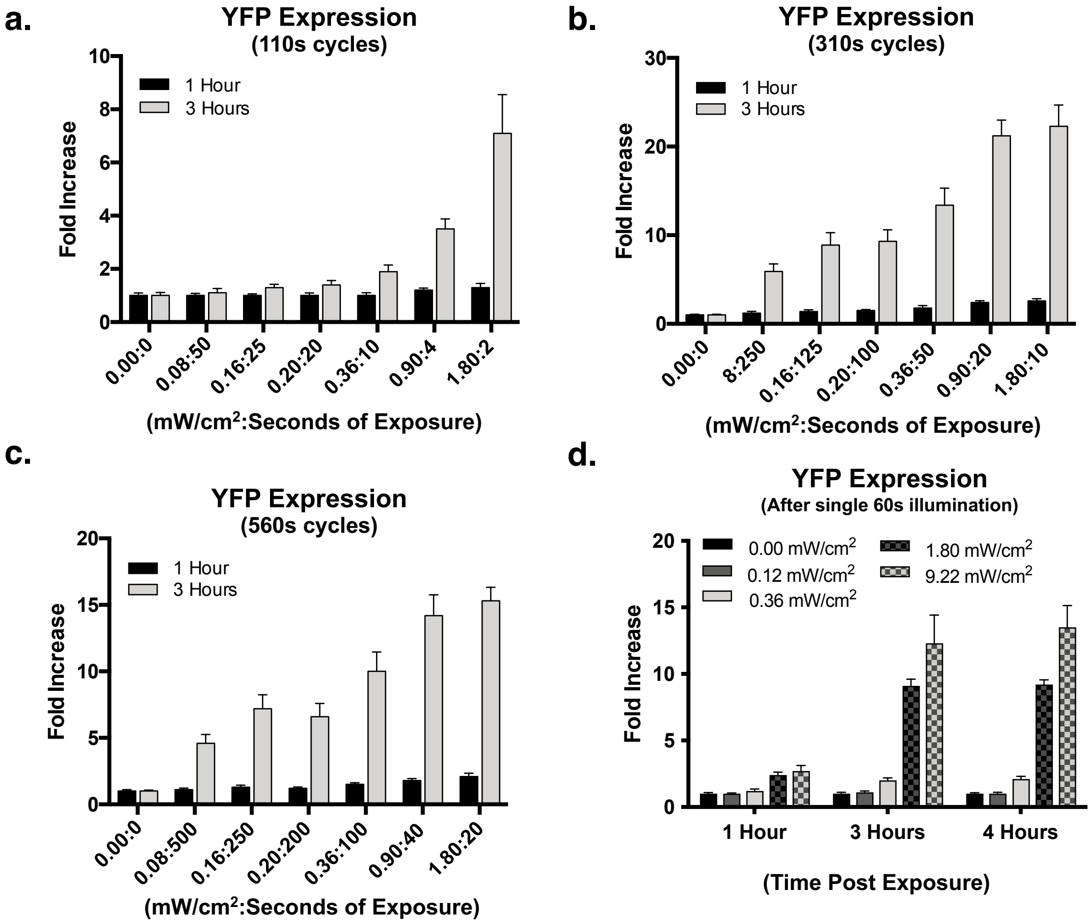


**Figure S4.** Blue light activated VP-EL222 driven YFP expression under various conditions. (a-c) YFP expression based on total duty cycle length (seconds of exposure + seconds off = total cycle length). For example, in panel a, sample 0.08:50 was exposed to 0.08 mW/cm$^2$ for 50s, and then in darkness for 60s totaling a 110s duty cycle. In this same panel, sample 1.80:2 was exposed to 1.80 mW/cm$^2$ for 2s, and then in darkness for 108s also totaling a 110s duty cycle. Thus, while light intensity changes, all samples in panel a are eventually exposed to the same number of photons by the end of the duty cycle. This approach is repeated in panels b and c with increasing total exposure. (d) YFP expression up to 4 hours after a single 60 second exposure at increasing blue light intensity. *Note: each mW/cm$^2$ was rounded for ease of viewing.*

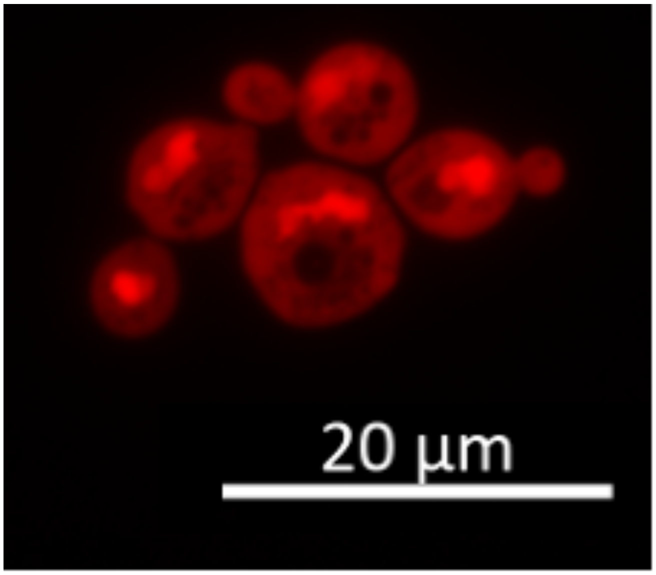


**Figure S5.** Confocal microscopy image of VP-mCherry-EL222 in *S. cerevisiae*.

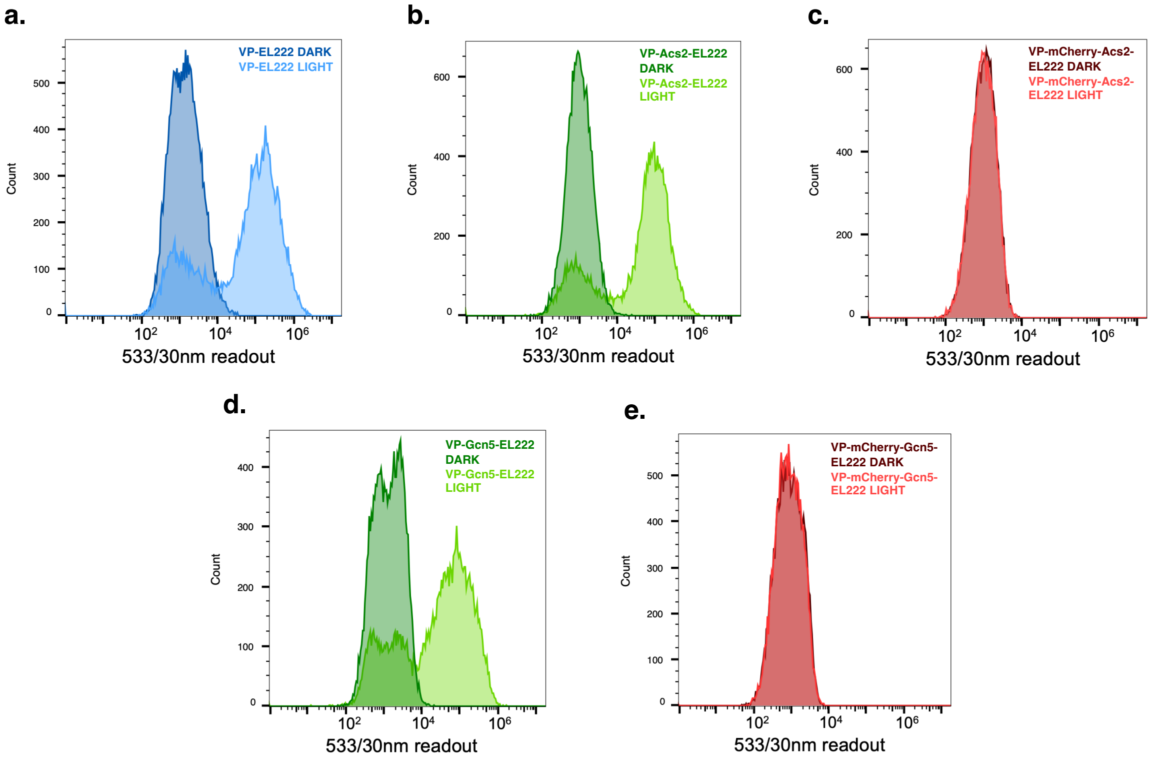


**Figure S6.** Flow cytometry histograms of various ACS2 and GCN5 fusion constructs after 4 hours of blue light exposure (20s on/60s off) with a light intensity of 0.90 mW/cm$^2$. (a) Original VP-EL222. (b) VP-Acs2-EL222. (c) VP-mCherry-Acs2-EL222. (d) VP-Gcn5-EL222. (e) VP-mCherry-Gcn5-EL222.

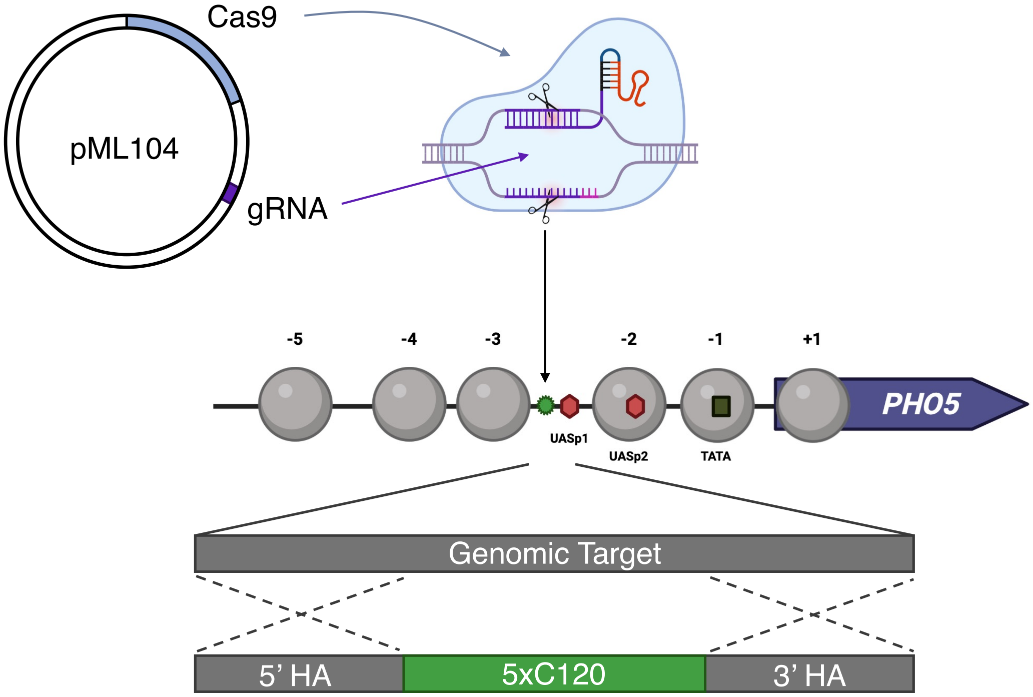


**Figure S7.** Outline of approach for stable 5xC120 genomic integration using CRISPR. The "Plug-and-Go" plasmid pML104 contains a constitutively active Cas9 protein as well as a gRNA producing sequence requiring only the 20bp target insertion via restriction enzyme ligation[3]. Electroporation of yeast cells with this plasmid as well as a knock-in cassette (5xC120 sequence flanked by 5' and 3' homology arms) leads to target integration. The Cas9 protein is loaded with the gRNA and produces a double stranded break. Yeast cells use double homologous recombination of the knock-in cassette which in turn alters the gRNA/PAM site preventing further Cas9 cutting.

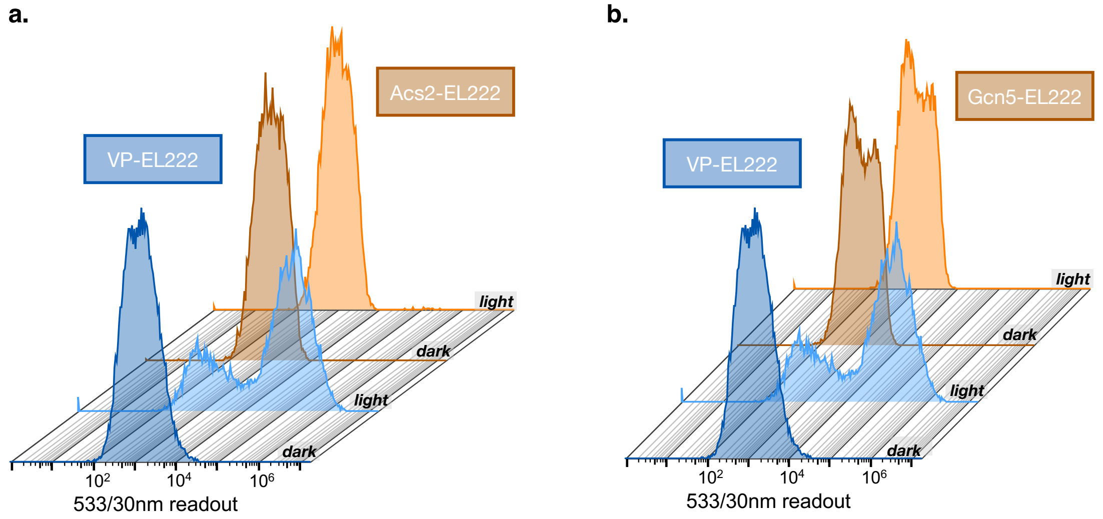


**Figure S8.** Flow cytometry histograms of Acs2 and Gcn5 EL222 fusion constructs in the Y422 plasmid lacking the VP16 transactivation domain after 4 hours of blue light exposure (20s on/60s off) with a light intensity of 0.90 mW/cm$^2$. (a) Waterfall plots showing the original VP-EL222 driving YFP production in the light. Acs2-EL222 lacking VP16 no longer drives YFP expression through direct Pol II recruitment. (b) Waterfall plots with a repeat of the original VP-EL222 driving YFP production in the light. Gcn5-EL222 also lacking VP16 no longer drives YFP expression in the same fashion.

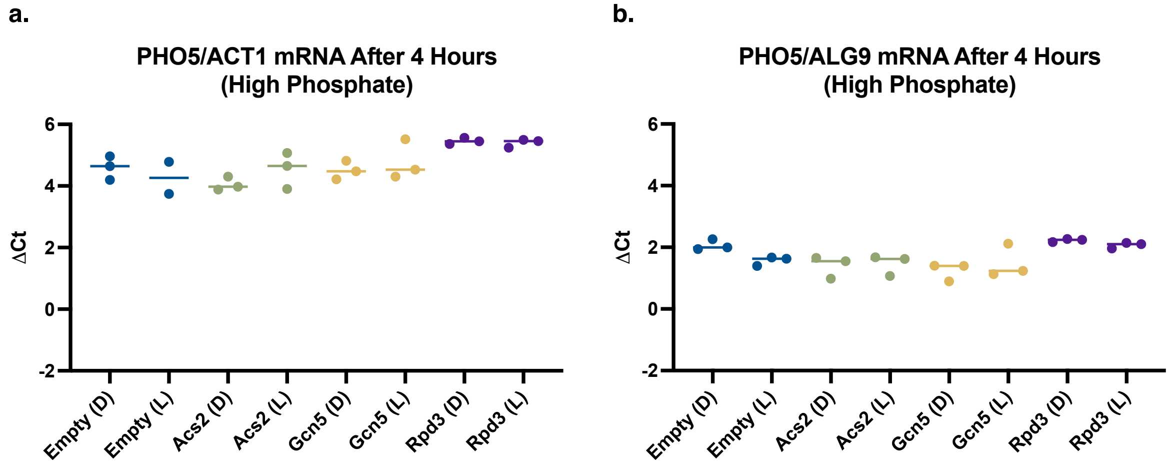


**Figure S9.** EL222 fusion constructs at the *PHO5* promoter in high $P_i$ (7.3 mM). qPCR runs of up to three technical replicates are plotted based on cycle thresholds. (a) *PHO5* transcripts in relation to the housekeeping gene *ACT1*. (b) *PHO5* transcripts in relation to the housekeeping gene *ALG9*. *n = 1*.

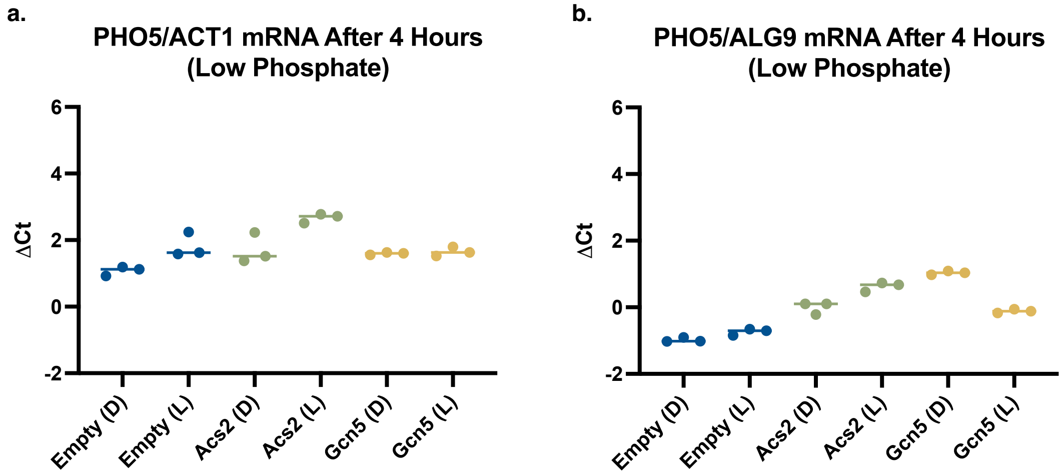


**Figure S10.** EL222 fusion constructs at the *PHO5* promoter in low $P_i$ (50 μM). qPCR runs of three technical replicates are plotted based on cycle thresholds. (a) *PHO5* transcripts in relation to the housekeeping gene *ACT1*. (b) *PHO5* transcripts in relation to the housekeeping gene *ALG9*. *n = 1*.

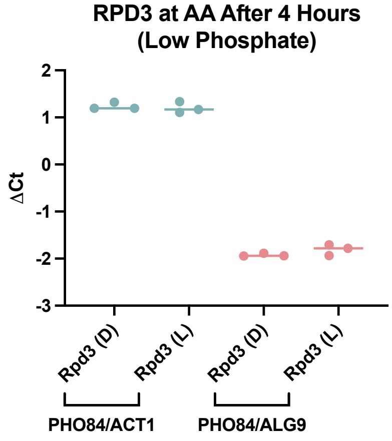


**Figure S11.** Rpd3-EL222 fusion constructs at the *PHO84* promoter in low $P_i$ (50 μM). qPCR runs of three technical replicates are plotted based on cycle thresholds. *PHO84* transcripts in relation to the housekeeping genes *ACT1* and *ALG9*. *n = 1*.

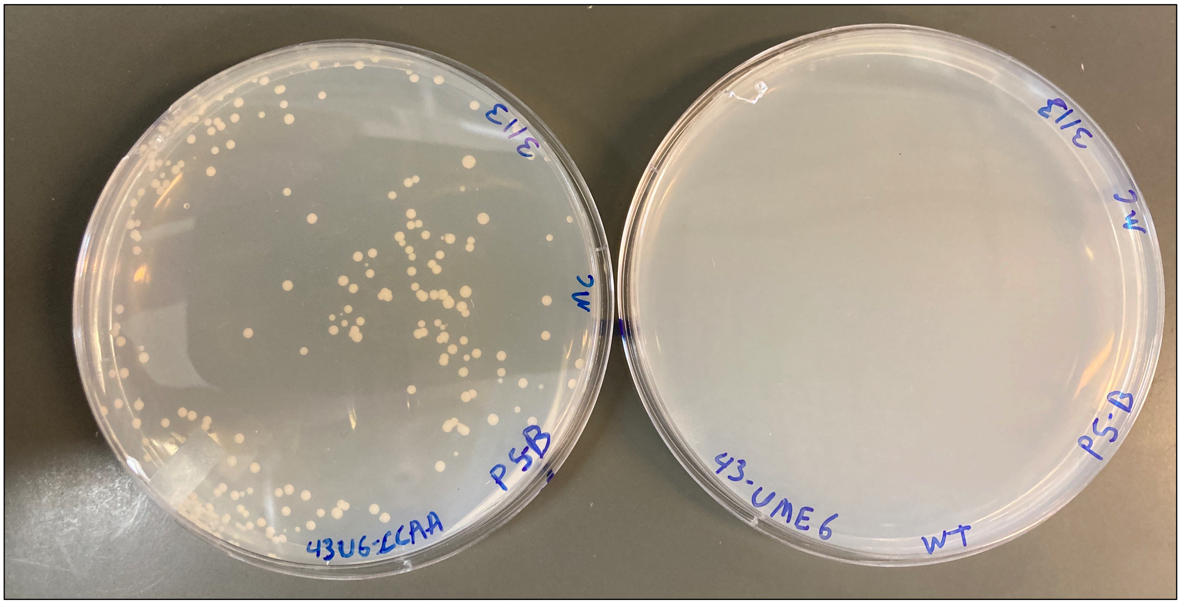


**Figure S12.** Lethality of wild type Ume6 protein as shown by comparison of Ume6-EL222 (right plates vs Ume6$^{C770A/C787A}$-EL222 (left plate) post transformation.

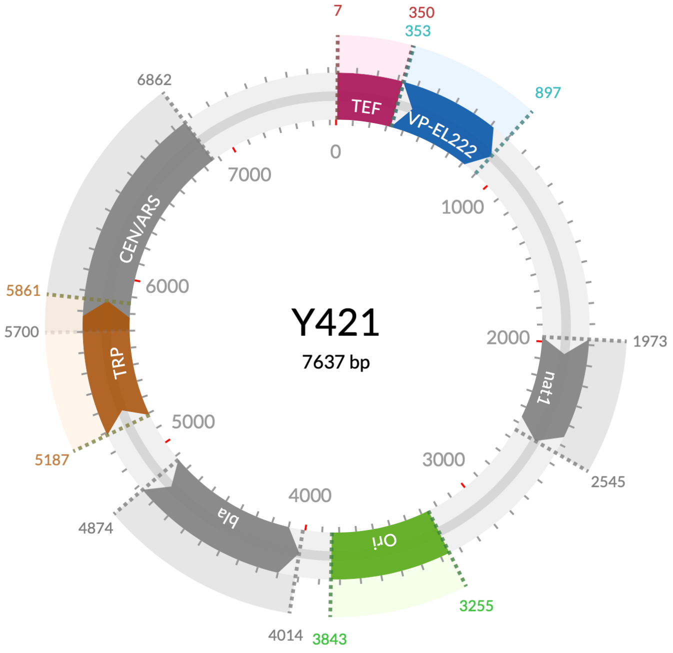


**Figure S13.** Map of the yeast centromeric plasmid Y421 consisting of a constitutively active VP-EL222 (blue) driven by an *Ashbya gossypii* TEF promoter (maroon), origin of replication (green), tryptophan auxotrophic marker (brown), antibiotic resistance markers beta-lactamase and nourseothricin (gray), and the *S. cerevisiae* CEN4 centromere fused to the autonomously replicating sequence ARS1 (gray).

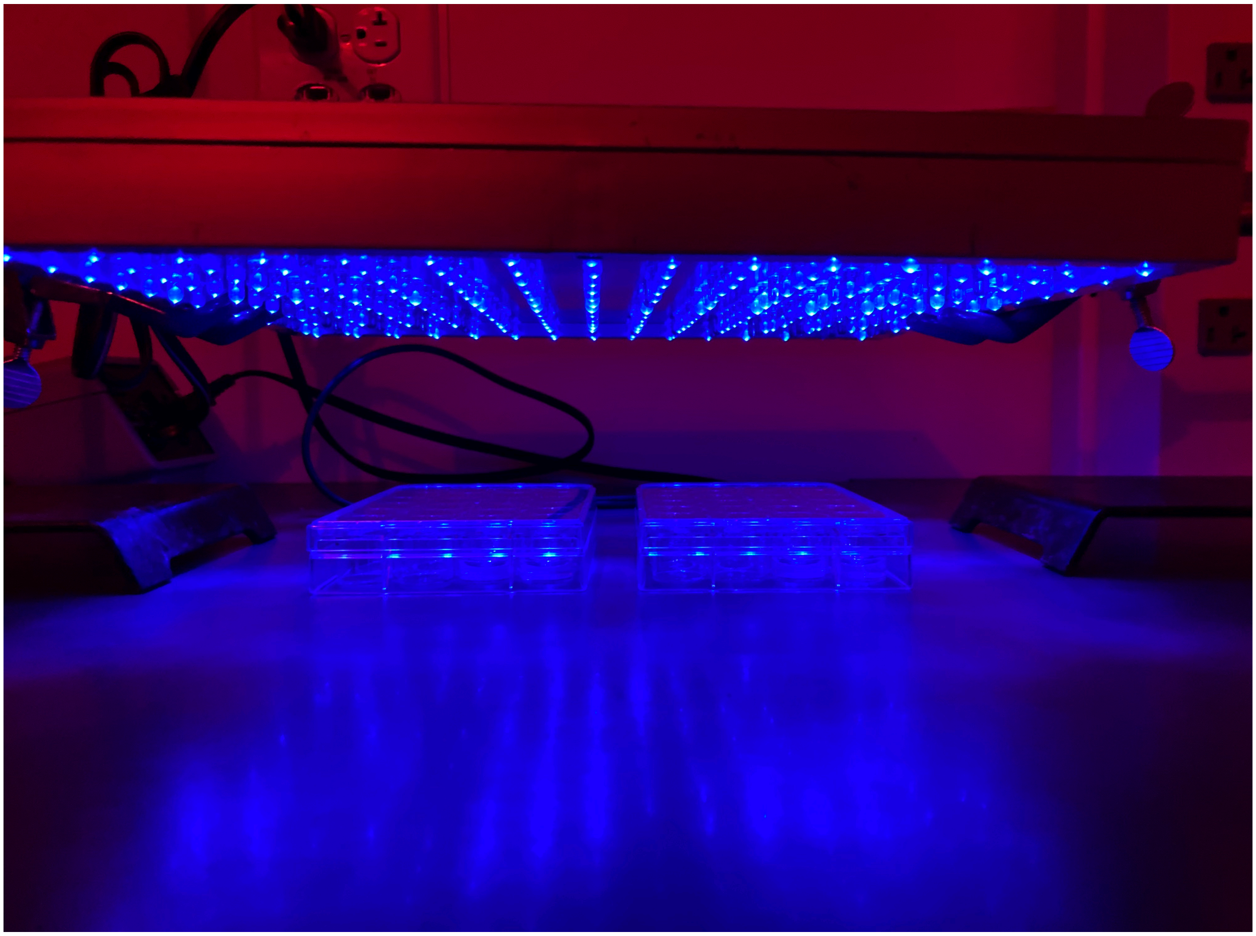

**Figure S14.** Overhead LED panel set up for phosphate based experiments. Cells were distributed in 24-well plates and exposed to repeating cycles of blue light (20s on/60s off) for 4 hours. The blue LED panel (465 nm, 13.8 Watts; 2501BU - LED Wholesalers) is positioned ~6 inches from the top of the plates and has an output of 15 $mW/cm^2$.

| Oligo Name | Sequence 5' to 3' | Notes |
|---|---|---|
| EL222-1F | **CAGACGACACACGCGTTGAG** | EL222 sequencing primer |
| EL222-1R | **TAGATTCCGGCTTCGACGGC** | EL222 sequencing primer |
| EL222-2F | **CGGGCTGGTGATGGAAAAGC** | EL222 sequencing primer |
| EL222-2R | **GACACGACCGATGCGATCG** | EL222 sequencing primer |
| 422-Bsp-NT-F | **GGGTCCGGAGGGGCAGACGACACACGCGTTGAGGTG** | Inserting BspEI RE into Y422 |
| 422-Bsp-NT-R | **TGCCCCTCCGGACCCACCGTACTCGTCAATTCCAAGGGC** | Inserting BspEI RE into Y422 |
| EL222-3F | **GGAGAAAACCGTCAAGATGC** | EL222 sequencing primer |
| EL222-3R | **CTTCTTCGGAATAGCCGGTC** | EL222 sequencing primer |
| Flat-NT-F | **CTTAAGCTAGGAGCATCCGGAGGGGCAGACGACAC** | Insert AflII into Y422 |
| Flat-NT-R | **TGCTCCTAGCTTAAGCCCACCGTACTCGTCAATTCCAAGG** | Insert AflII into Y422 |
| mCAflNCTex-F | **TAAGCACTTAAGGGTGGTGGATCAGGTGGAG** | AflII Overhangs with additional 6bp |
| mCBspNTex-R | **TGCTTATCCGGACTTGTACAGCTCGTCCATGCC** | BspEI Overhangs with additional 6bp |
| GS3B1-T | **CCGGAGGTGGCGGTGGCTCGGGCGGAGGTGGGTCGGGTGG CGGCGGATCAT** | 5' phosphorylated (GGGGS)3 linker w/ BspEI RE |
| GS3B1-B | **CCGGATGATCCGCCGCCACCCGACCCACCTCCGCCCGAGCC ACCGCCACCT** | 5' phosphorylated (GGGGS)3 linker w/ BspEI RE |
| GS3B2-T | **CCGGAGGTGGAGGAGGCTCTGGTGGAGGCGGTAGCGGAGG CGGAGGGTCGT** | 5' phosphorylated (GGGGS)3 linker w/ BspEI RE |
| GS3B2-B | **CCGGACGACCCTCCGCCTCCGCTACCGCCTCCACCAGAGCCT CCTCCACCT** | 5' phosphorylated (GGGGS)3 linker w/ BspEI RE |
| NsiINT2-F | **GACGAGCTGTACAAGATGCATCTAGGAGCATCCGGAGGGGC AGACGAC** | Insert NsiI RE + scaffold (39R partner) |
| NTlink2-R | **CTTGTACAGCTCGTCCATGCCGCCGGTGGAGTG** | Linker primer between mCherry/EL222 |
| DEcoRI-F | **CCAGTGAGTTCGACATGGAGGCCCAGAATACCCTCC** | Mutate EcoRI to nothing to free up EcoRI |
| DEcoRI-R | **ATGTCGAACTCACTGGCCGTCGTTTTACAACGTCGTGAC** | Mutate EcoRI to nothing to free up EcoRI |
| EcoSeq-F | **CCAGCGATATATGCGGTGTG** | Covers EcoRI region for Sequence Check |
| YFP-Seq-F | **CCAGTTCCATGGCCAACCTTAG** | Covers YFP Sequence in Y422 |
| ASC2-EcoRI-F | **TAAGCAGAATTCACAATCAAGGAACATAAAGTAGTTTATG** | PCR ACS2 for insertion into mCherry scaffold |
| ASC2-XbaI-R | **TGCTTATCTAGATTTCTTTTTTTGAGAGAAAAATTGGTTC** | PCR ACS2 for insertion into mCherry scaffold |
| mC222-1F | **GCATGGACGAGCTGTACAAG** | Sequencing check/+112R to make NLS_GCN5 |
| mC222-1R | **CTCAACGCGTGTGTCGTCTG** | Sequencing check |
| Aat-Sand-T | **TAAGCAGAATTCCTTGGAGCACTTGACGTCGGAGCACTTGCA TCTAGAGCTATC** | AatII scaffold self-anneal |
| Aat-Sand-B | **GATAGCTCTAGATGCAAGTGCTCCGACGTCAAGTGCTCCAAG GAATTCTGCTTA** | AatII scaffold self-anneal |
| GCN5-AatII-F | **TAAGCAGACGTCGTCACAAAACATCAGATTGAAGAG** | PCR GCN5 for insertion into 42mCscaff |
| GCN5-XbaI-R | **TGCTTATCTAGAATCAATAAGGTGAGAATATTCAGG** | PCR GCN5 for insertion into 42mCscaff |
| GCN5seq-F | **GTCACAAAACATCAGATTGAAGAGGATCAC** | Sequence GCN5 insertion |
| GCN5seq-R | **GGTCTCAGCCTGCTCAGGTTGTATTTC** | Sequence GCN5 insertion |
| NVEco-F | **GGATCAGAATTCACAATCAAGGAAC** | EcoRI deletion of mCherry and Linker ACS2 |
| NVEco-Ra | **TGACTCGAATTCCCCACCGTACTCGTCAATTC** | EcoRI deletion of mCherry and Linker |

| | | |
|---|---|---|
| NVG5-At-Rb | **TGACTCGACGTCCCCACCGTACTCGTCAATTCC** | AatII deletion of mCherry and Linker (72F Partner) |
| VpEco-R2 | **TGCTTAGAATTCGACTTTACGCTTCTTTTTAGGGCC** | Delete VP16 and Insert EcoRI |
| NG5-At-R | **TGACTCGACGTCGACTTTACGCTTCTTTTTAGGGCCC** | Delete VP16/mCherry and insert AatII |
| 43NotI-F | **TAAGCAGCGGCCGCCTGGATCCACAGTTTATTCCTGG** | Turn 422 into 423 using NotI (Removes 5xC120 + YFP) |
| 43NotI-R | **TGCTTAGCGGCCGCGTTATCCCTAGCGGATCTGC** | Turn 422 into 423 using NotI (Removes 5xC120 + YFP) |
| ConAatII-F | **TAAGCAGACGTCGGGGCAGACGACACACGC** | EL222-GCN5 into EL222 Without TAD |
| ConAatII-R | **TGTGACGACGTCGACTTTACGCTTC** | EL222-GCN5 into EL222 Without TAD |
| Pho4-Eco-F | **TAAGCAGAATTCGGCCGTACAACTTCTGAGGG** | PHO4 into 423 making fusion construct |
| Pho4-Spe-R | **TGCTTAACTAGTCGTGCTCACGTTCTGCTGTAG** | PHO4 into 423 making fusion construct |
| P4-E259L-F | **CATGCACTGCAAGCACGGCGTAATCGATTAGCGGTC** | PHO4 mutation (no DNA binding) |
| P4-E259L-R | **TGCTTGCAGTGCATGCTTGTGTGATTCGCGCTTGTC** | PHO4 mutation (no DNA binding) |
| P4-Seq1-F | **GTCGTAGCAAGTGAGTCTCC** | PHO4 fusion sequencing primer |
| ACS2-T325K-F2 | **CTGGATCAAGGGTCACACCTATGCTCTATATGGTCCATTAACC TTG** | ACS2 catalytically dead mutant |
| ACS2-T325K-R | **TGACCCTTGATCCAGCCGACGTCACCGGCAGTGAAG** | ACS2 catalytically dead mutant |
| ACS2-Seq1-F | **GCAGAGAACTTACCTACCTC** | ACS2 catalytically dead sequencing primer |
| GC5-E173A-F | **TTCGCAGCAATTGTTTTCTGTGCCATCAGTTCGTCG** | GCN5 catalytically dead mutant |
| GC5-E173A-R | **AACAATTGCTGCGAATTCTCTCTTATCGAAAGGTCGATATG** | GCN5 catalytically dead mutant |
| GC5-Seq1-F | **GAGACCCAGTGTCGTAGAGG** | GCN5 catalytically dead sequencing primer |
| RPD3-F | **TAAGCACAATTGGTATATGAAGCAACACCTTTTGATCC** | RPD3 into 423 making fusion construct |
| RPD3-R | **TGCTTATCTAGAATAGAATTCATTGTCATGCTCAACATG** | RPD3 into 423 making fusion construct |
| UME6-F1 | **TAAGCACAATTGCTAGACAAGGCGCGCTCTC** | UME6 into 423 making fusion construct |
| UME6-R1 | **TGCTTATCTAGATTTTTTTTTCATTGCTCTTCTTTTGGCCTC** | UME6 into 423 making fusion construct |
| U6C770A-F2 | **ACTGGTGCCTGGATTTGTAGATTAAGGAAAAAGAAGTG** | UME6 C770A mutation primers |
| U6C770A-R2 | **ATCCAGGCACCAGTACGGGACCTTGTACCTTG** | UME6 C770A mutation primers |
| U6C787A-F1 | **CGCACGCTTTCAACTGTGAAAGGTTGAAATTGGAC** | UME6 C787A mutation primers |
| U6C787A-R1 | **GTTGAAAGCGTGCGGTCTTTCCTCGGTACAC** | UME6 C787A mutation primers |
| U6mut-Seq-F | **CTTGGCGAATCCTCAACTTC** | Sequence confirmation of UME6 mutations |
| P5g2Crsp-T | **GATCAAACTTCAAACGAAGGTAAAGTTTTAGAGCTAG** | gRNA for pML104 PHO5 - B |
| P5g2Crsp-B | **CTAGCTCTAAAACTTTACCTTCGTTTGAAGTTT** | gRNA for pML104 PHO5 - B |
| gRNA-C-T | **GATCACGCTCTCTTTACAGGACGCGTTTTAGAGCTAG** | gRNA for pML104 PHO5 - C |
| gRNA-C-B | **CTAGCTCTAAAACGCGTCCTGTAAAGAGAGCGT** | gRNA for pML104 PHO5 - C |
| gRNA-E-T | **GATCTGTCAGAGATCAAGTCAGAGGTTTTAGAGCTAG** | gRNA for pML104 PHO5 - E |
| gRNA-E-B | **CTAGCTCTAAAACCTCTGACTTGATCTCTGACA** | gRNA for pML104 PHO5 - E |
| gRNA-F-T | **GATCGTCTACAACACCGTCTTCCAGTTTTAGAGCTAG** | gRNA for pML104 PHO5 - F |

| | | |
|---|---|---|
| gRNA-F-B | **CTAGCTCTAAAACTGGAAGACGGTGTTGTAGAC** | gRNA for pML104 PHO5 - F |
| gRNA-AA-T | **GATCTAATAATTAGCTCATAGATGGTTTTAGAGCTAG** | gRNA for pML104 PHO84 |
| gRNA-AA-B | **CTAGCTCTAAAACCATCTATGAGCTAATTATTA** | gRNA for pML104 PHO84 |
| gRNA-BB-T | **GATCGCTATGAAGAAAATGTTGCCGTTTTAGAGCTAG** | gRNA for pML104 PHO84 |
| gRNA-BB-B | **CTAGCTCTAAAAC GGCAACATTTTCTTCATAGC** | gRNA for pML104 PHO84 |
| C120-F | **GCTAGCCTCGAGTAGGTAGC** | XhoI-5xC120 for cassettes |
| C120-R | **CCTCTAGTGTCTAAGCTTCATGG** | HindIII-5xC120 for cassettes |
| P5g2-5'F | **TAAGCAGAATTCTATGTTCTATTTACTGACCGAAAGTAGC** | 5' Arm PHO5 gRNA B |
| P5g2-5'R | **TAAGCACTCGAGTACCTTCGTTTGAAGTTTATAGACGAC** | 5' Arm PHO5 gRNA B |
| P5g2-3'F | **TGCTTAAAGCTTTTCATAGCGCTTTTTCTTTGTCTGC** | 3' Arm PHO5 gRNA B |
| P5g2-3'R | **TGCTTAACTAGTAAGTAAGGTGACCAATTTGATAATTTGG** | 3' Arm PHO5 gRNA B |
| Pho5CAS-F | **CTGACCGAAAGTAGCTCGCTAC** | PCR Full B' cassette |
| Pho5CAS-R | **TGGCATGTGCGATCTCTTCG** | PCR Full B' cassette |
| C5'EcoRI-F1 | **TAAGCAGAATTCCTGCACATTGGCATTAGCTAGG** | 5' Arm PHO5 gRNA C |
| C5'XhoI-R1 | **TAAGCACTCGAGGTCCTGTAAAGAGAGCGTGC** | 5' Arm PHO5 gRNA C |
| C3'HindIII-F1 | **TGCTTAAAGCTTAGACCGGCATTACAAGGATCC** | 3' Arm PHO5 gRNA C |
| C3'SpeI-R1 | **TGCTTAACTAGTGGCATTGTGCAGTTGCGTTC** | 3' Arm PHO5 gRNA C |
| E5'EcoRI-F1 | **TAAGCAGAATTCCCAGCAAGATGAGCCTTACC** | 5' Arm PHO5 gRNA E |
| E5'XhoI-R1 | **TAAGCACTCGAGCTGACTTGATCTCTGACATTAAAAATTCG** | 5' Arm PHO5 gRNA E |
| E3'HindIII-F1 | **TGCTTAAAGCTTCTGCCTTTACCGTAATTTTCAATTGC** | 3' Arm PHO5 gRNA E |
| E3'SpeI-R1 | **TGCTTAACTAGTCTAATGCCAATGTGCAGTAGTAAC** | 3' Arm PHO5 gRNA E |
| F5'EcoRI-F1 | **TAAGCAGAATTCGCCAATAGAATAAGAGGTAGG** | 5' Arm PHO5 gRNA F |
| F5'XhoI-R1 | **TAAGCACTCGAGTATTTTCAATTTCTTCCTCAAAATTTG** | 5' Arm PHO5 gRNA F |
| F3'HindIII-F1 | **TGCTTAAAGCTTGAAGACGGTGTTGTAGACGATAATG** | 3' Arm PHO5 gRNA F |
| F3'SpeI-R1 | **TGCTTAACTAGTCTGTCAAATACTGTGTTACCCCC** | 3' Arm PHO5 gRNA F |
| AA5'EcoRI-F1 | **TAAGCAGAATTCCGGCTATGAAGAAAATGTTGCCC** | 5' Arm PHO5 gRNA AA |
| AA5'XhoI-R1 | **TAAGCACTCGAGTCGGTTAATTAATGAGTAATACGCACG** | 5' Arm PHO5 gRNA AA |
| AA3'HindIII-F1 | **TGCTTAAAGCTTTCTATGAGCTAATTATTATTCCTTTTTGGC** | 3' Arm PHO5 gRNA AA |
| AA3'SpeI-R1 | **TGCTTAACTAGTGACGGAACTCATTTGGATTGTATTC** | 3' Arm PHO5 gRNA AA |
| BB5'EcoRI-F1 | **TAAGCAGAATTCCAGCCGCTCGTTTATCAACC** | 5' Arm PHO5 gRNA BB |
| BB5'XhoI-R1 | **TAAGCACTCGAGCAACATTTTCTTCATAGCCGCTAG** | 5' Arm PHO5 gRNA BB |
| BB3'HindIII-F1 | **TGCTTAAAGCTTCTGAAAAACACCCGTTCCTC** | 3' Arm PHO5 gRNA BB |
| BB3'SpeI-R1 | **TGCTTAACTAGTCTAGCTAATAAGCAGGCAAAACG** | 3' Arm PHO5 gRNA BB |
| C'-Cas-F | **CTGCACATTGGCATTAGCTAGG** | PCR Full C' cassette |
| C'-Cas-R | **GGCATTGTGCAGTTGCGTTC** | PCR Full C' cassette |
| E'-Cas-F | **CCAGCAAGATGAGCCTTACC** | PCR Full E' cassette |
| E'-Cas-R | **CTAATGCCAATGTGCAGTAGTAAC** | PCR Full E' cassette |
| F'-Cas-F | **GCCAATAGAATAAGAGGTAGG** | PCR Full F' cassette |
| F'-Cas-R | **CTGTCAAATACTGTGTTACCC** | PCR Full F' cassette |

| | | |
|---|---|---|
| AA'-Cas-F | **CGGCTATGAAGAAAATGTTGCCC** | PCR Full AA' cassette |
| AA'-Cas-R | **GACGGAACTCATTTGGATTGTATTCG** | PCR Full AA' cassette |
| BB'-Cas-F | **CAGCCGCTCGTTTATCAACC** | PCR Full BB' cassette |
| BB'-Cas-R | **CTAGCTAATAAGCAGGCAAAACGG** | PCR Full BB' cassette |
| PosPCR-F1 | **GTCAGTCCCCACGCTAATAG** | Sequence inserts |
| PosPCR-F2 | **GAGACTCCGTCCCTCTTTAG** | Sequence inserts |
| PosPCR-R1 | **GCAGTGTCATGAACCCTTTC** | Sequence inserts |
| PosPCR-R2 | **CACCCAGGAATGGGAATATG** | Sequence inserts |
| C'seq-F | **GAGACTCCGTCCCTCTTTAGTG** | Sequence inserts |
| C'seq-R | **GTGAGTGCCAAGGTTGTATCG** | Sequence inserts |
| E'seq-F | **CCTAATTTGTGGCTGGATAAATGG** | Sequence inserts |
| E'seq-R | **CGGACCTACAACAAAGTTGGG** | Sequence inserts |
| F'seq-R | **GATCAACGCTTTCCCAGACTTC** | Sequence inserts |
| AA'seq-F | **GCACGTTGGTGCTGTTATAGG** | Sequence inserts |
| AA'seq-R | **GTGGAAGGCCATGTTACCAC** | Sequence inserts |
| BB'seq-F | **ACGACTCGGTATACTCTGCC** | Sequence inserts |
| BB'seq-R | **CATTACCGGTGTGCAATGTG** | Sequence inserts |
| qPCR-P5-F1 | **TACCCTACTGTCAGTCTGGC** | qPCR target for PHO5 transcripts |
| qPCR-P5-R1 | **GGCGTTCATTTCACCAGTGT** | qPCR target for PHO5 transcripts |
| qPCR-ACT1-F | **CGAATTGAGAGTTGCCCCAG** | qPCR target for ACT1 HKG transcripts |
| qPCR-ACT1-R | **CAAGGACAAAACGGCTTGGA** | qPCR target for ACT1 HKG transcripts |
| qPCR-ALG9-F | **ACGGATAGTGGCTTTGGTGA** | qPCR target for ALG9 HKG transcripts |
| qPCR-ALG9-R | **TCTGGCAGCAGGAAAGAACT** | qPCR target for ALG9 HKG transcripts |
| qPCR-P84-F | **GACCGCTTTGTTCTGTGTCA** | qPCR target for PHO84 transcripts |
| qPCR-P84-R | **GCGGTTTGTGCAATAATGGC** | qPCR target for PHO84 transcripts |

**Table S1.** List of primers used in this study.
