## Supplemental Sequence File S1 for "Optogenetic control of phosphate-responsive genes using single component fusion proteins in *Saccharomyces cerevisiae*"

**Y422 Plasmid**

CEN/CEN

NLS

VP16

EL222

5xC120

YFP

nat1

Ori

bla

TRP/TRP

agcttacattttatgttagctggtggactgacgccagaaaatgttggtgatgcgcttagattaaatggcgttattggtgttgatgtaagcggaggtgtggagacaaatggtgtaaaagactctaacaaaatagcaaatttcgtcaaaaatgctaagaaataggttattactgagtagtatttatttaagtattgtttgtgcacttgcctgcaagccttttgaaaagcaagcataaaagatctaaacataaaatctgtaaaataacaagatgtaaagataatgctaaatcatttggctttttgattgattgtacaggaaaatatacatcgcagggggttgacttttaccatttcaccgcaatggaatcaaacttgttgaagagaatgttcacaggcgcatacgctacaatgacccgattcttgctagccttttctcggtcttgcaaacaaccgccggcagcttagtatataaatacacatgtacatacctctctccgtatcctcgtaatcattttcttgtatttatcgtcttttcgctgtaaaaactttatcacacttatctcaaatacacttattaaccgcttttactattatcttctacgctgacagtaatatcaaacagtgacacatattaaacacagtggtttctttgcataaacaccatcagcctcaagtcgtcaagtaaagatttcgtgttcatgcagatagataacaatctatatgttgataattagcgttgcctcatcaatgcgagatccgtttaaccggaccctagtgcacttaccccacgttcggtccactgtgtgccgaacatgctccttcactattttaacatgtggaattaattctcatgtttgacagcttatcatcgaactctaagaggtgatacttatttactgtaaaactgtgacgataaaaccggaaggaagaataagaaaactcgaactgatctataatgcctattttctgtaaagagtttaagctatgaaagcctcggcattttggccgctcctaggtagtgctttttttccaaggacaaaacagtttctttttcttgagcaggttttatgtttcggtaatcataaacaataaataaattatttcatttatgtttaaaaataaaaaataaaaaagtattttaaatttttaaaaaagttgattataagcatgtgaccttttgcaagcaattaaattttgcaatttgtgattttaggcaaaagttacaatttctggctcgtgtaatatatgtatgctaaagtgaacttttacaaagtcgatatggacttagtcaaaagaaattttcttaaaaatatatagcactagccaatttagcacttctttatgagatatattatagactttattaagccagatttgtgtattatatgtatttacccggcgaatcatggacatacattctgaaataggtaatattctctatggtgagacagcatagataacctaggatacaagttaaaagctagtactgttttgcagtaatttttttcttttttataagaatgttaccacctaaataagttataaagtcaatagttaagtttgatatttgattgtaaaataccgtaatatatttgcatgatcaaaaggctcaatgttgactagccagcatgtcaaccactatattgatcaccgatatatggacttccacaccaactagtaatatgacaataaattcaagatattcttcatgagaatggcccagcgatatatgcggtgtgaaataccgcacagatgcgtaaggagaaaataccgcatcaggcgccattcgccattcaggctgcgcaactgttgggaagggcgatcggtgcgggcctcttcgctattacgccagctggcgaaagggggatgtgctgcaaggcgattaagttgggtaacgccagggttttcccagtcacgacgttgtaaaacgacggccagtgaattcgacatggaggcccagaataccctccttgacagtcttgacgtgcgcagctcaggggcatgatgtgactgtcgcccgtacatttagcccatacatccccatgtataatcatttgcatccatacattttgatggccgcacggcgcgaagcaaaaattacggctcctcgctgcagacctgcgagcagggaaacgctcccctcacagacgcgttgaattgtccccacgccgcgcccctgtagagaaatataaaaggttaggatttgccactgaggttcttctttcatatacttccttttaaaatcttgctaggatacagttctcacatcacatccgaacataaacaaccgcatgggccctaaaaagaagcgtaaagtcgcccccccgaccgatgtcagcctgggggacgagctccacttagacggcgaggacgtggcgatggcgcatgccgacgcgctagacgatttcgatctggacatgttgggggacggggattccccgggtccgggatttaccccccacgactccgccccctacggcgctctggatatggccgacttcgagtttgagcagatgtttaccgatgcccttggaattgacgagtacggtggggaattcggggcagacgacacacgcgttgaggtgcaaccgccggcgcagtgggtcctcgacctgatcgaggccagcccgatcgcatcggtcgtgtccgatccgcgtctcgccgacaatccgctgatcgccatcaaccaggccttcaccgacctgaccggctattccgaagaagaatgcgtcggccgcaattgccgattcctggcaggttccggcaccgagccgtggctgaccgacaagatccgccaaggcgtgcgcgagcacaagccggtgctggtcgagatcctgaactacaagaaggacggcacgccgttccgcaatgccgtgctcgttgcaccgatctacgatgacgacgacgagcttctctatttcctcggcagccaggtcgaagtcgacgacgaccagcccaacatgggcatggcgcgccgcgaacgcgccgcggaaatgctcaagacgctgtcgccgcgccagctcgaggttacgacgctggtggcatcgggcttgcgcaacaaggaagtggcggcccggctcggcctgtcggagaaaaccgtcaagatgcaccgcgggctggtgatggaaaagctcaacctgaagaccagtgccgatctggtgcgcattgccgtcgaagccggaatctaaccacttctaaataagcgaatttcttatgatttatgatttttattattaaataagttataaaaaaaataagtgtatacaaattttaaagtgactcttaggttttaaaacgaaaattcttattcttgagtaactctttcctgtaggtcaggttgctttctcaggtatagtatgaggtcgctcttattgaccacacctctaccggcagatccgctagggataacgtcgctagcctcgagtaggtagcctttagtccatgcgttataggtagcctttagtccatgcgttataggtagcctttagtccatgcgttataggtagcctttagtccatgcgttataggtagcctttagtccatgaagcttagacactagagggtatataatggaagctcgacttccagcttggcaatccggtactgttggtaaaggatccactagtccagtgtggtgaagcttttaattaataacaaaatgtctaaaggtgaagaattattcactggtgttgtcccaactttggttgaattagatggtgatgttaatggtcacaaattttctgtctccggtgaaggtgaaggtgatgctacttacggtaaattgaccttaaaattaatttgtactactggtaaattgccagttccatggccaaccttagtcactactttaggttatggtttaatgtgttttgctagatacccagatcatatgaaacaacatgactttttcaagtctgccatgccagaaggttatgttcaagaaagaactatttttttcaaagatgacggtaactacaagaccagagctgaagtcaagtttgaaggtgataccttagttaatagaatcgaattaaaaggtattgattttaaagaagatggtaacattttaggtcacaaattggaatacaactataactctcacaatgtttacatcactgctgacaaacaaaagaatggtatcaaagctaacttcaaaattagacacaacattgaagatggtggtgttcaattagctgaccattatcaacaaaatactccaattggtgatggtccagtcttgttaccagacaaccattacttatcctatcaatctgccttatccaaagatccaaacgaaaagagagaccacatggtcttgttagaatttgttactgctgctggtattacccatggtatggatgaattgtacaaataaggcgcgccactgacaataaaaagattcttgttttcaagaacttgtcatttgtatagtttttttatattgtagttgttctattttaatcaaatgttagcgtgatttatattttttttcgcctcgacatcatctgcccagatgcgaagttaagtgcgcagaaagtaatatcatgcgtcaatcgtatgtgaatgctggtcgctatactggatccacagtttattcctggcatccactaaatataatggagcccgctttttaagctggcatccagaaaaaaaaagaatcccagcaccaaaatattgttttcttcaccaaccatcagttcataggtccattctcttagcgcaactacagagaacaggggcacaaacaggcaaaaaacgggcacaacctcaatggagtgatgcaacctgcctggagtaaatgatgacacaaggcaattgacccacgcatgtatctatctcattttcttacaccttctattaccttctgctctctctgatttggaaaaagctgaaaaaaaaggttgaaaccagttccctgaaattattcccctacttgactaataagtatataaagacggtaggtattgattgtaattctgtaaatctatttcttaaacttcttaaattctacttttatagttagtcttttttttagttttaaaacaccaagaacttagtttcgaataaacacacataaacaaaatgggtaccactcttgacgacacggcttaccggtaccgcaccagtgtcccgggggacgccgaggccatcgaggcactggatgggtccttcaccaccgacaccgtcttccgcgtcaccgccaccggggacggcttcaccctgcgggaggtgccggtggacccgcccctgaccaaggtgttccccgacgacgaatcggacgacgaatcggacgacggggaggacggcgacccggactcccggacgttcgtcgcgtacggggacgacggcgacctggcgggcttcgtggtcgtctcgtactccggctggaaccgccggctgaccgtcgaggacatcgaggtcgccccggagcaccgggggcacggggtcgggcgcgcgttgatggggctcgcgacggagttcgcccgcgagcggggcgccgggcacctctggctggaggtcaccaacgtcaacgcaccggcgatccacgcgtaccggcggatggggttcaccctctgcggcctggacaccgccctgtacgacggcaccgcctcggacggcgagcaggcgctctacatgagcatgccctgcccctaattaattaaggccgctagggccctgcaggagggccgcatcatgtaattagttatgtcacgcttacattcacgccctccccccacatccgctctaaccgaaaaggaaggagttagacaacctgaagtctaggtccctatttatttttttatagttatgttagtattaagaacgttatttatatttcaaatttttcttttttttctgtacagacgcgtgtacgcatgtaacattatactgaaaaccttgcttgagaaggttttgggacgctcgaaggctttaatttgcggccaagcttggcgtaatcatggtcatagctgtttcctgtgtgaaattgttatccgctcacaattccacacaacatacgagccggaagcataaagtgtaaagcctggggtgcctaatgagtgagctaactcacattaattgcgttgcgctcactgcccgctttccagtcgggaaacctgtcgtgccagctgcattaatgaatcggccaacgcgcggggagaggcggtttgcgtattgggcgctcttccgcttcctcgctcactgactcgctgcgctcggtcgttcggctgcggcgagcggtatcagctcactcaaaggcggtaatacggttatccacagaatcaggggataacgcaggaaagaacatgtgagcaaaaggccagcaaaaggccaggaaccgtaaaaaggccgcgttgctggcgtttttccataggctccgcccccctgacgagcatcacaaaaatcgacgctcaagtcagaggtggcgaaacccgacaggactataaagataccaggcgtttccccctggaagctccctcgtgcgctctcctgttccgaccctgccgcttaccggatacctgtccgcctttctcccttcgggaagcgtggcgctttctcatagctcacgctgtaggtatctcagttcggtgtaggtcgttcgctccaagctgggctgtgtgcacgaaccccccgttcagcccgaccgctgcgccttatccggtaactatcgtcttgagtccaacccggtaagacacgacttatcgccactggcagcagccactggtaacaggattagcagagcgaggtatgtaggcggtgctacagagttcttgaagtggtggcctaactacggctacactagaaggacagtatttggtatctgcgctctgctgaagccagttaccttcggaaaaagagttggtagctcttgatccggcaaacaaaccaccgctggtagcggtggtttttttgtttgcaagcagcagattacgcgcagaaaaaaaggatctcaagaagatcctttgatcttttctacggggtctgacgctcagtggaacgaaaactcacgttaagggattttggtcatgagattatcaaaaaggatcttcacctagatccttttaaattaaaaatgaagttttaaatcaatctaaagtatatatgagtaaacttggtctgacagttaccaatgcttaatcagtgaggcacctatctcagcgatctgtctatttcgttcatccatagttgcctgactccccgtcgtgtagataactacgatacgggagggcttaccatctggccccagtgctgcaatgataccgcgagacccacgctcaccggctccagatttatcagcaataaaccagccagccggaagggccgagcgcagaagtggtcctgcaactttatccgcctccatccagtctattaattgttgccgggaagctagagtaagtagttcgccagttaatagtttgcgcaacgttgttgccattgctacaggcatcgtggtgtcacgctcgtcgtttggtatggcttcattcagctccggttcccaacgatcaaggcgagttacatgatcccccatgttgtgcaaaaaagcggttagctccttcggtcctccgatcgttgtcagaagtaagttggccgcagtgttatcactcatggttatggcagcactgcataattctcttactgtcatgccatccgtaagatgcttttctgtgactggtgagtactcaaccaagtcattctgagaatagtgtatgcggcgaccgagttgctcttgcccggcgtcaatacgggataataccgcgccacatagcagaactttaaaagtgctcatcattggaaaacgttcttcggggcgaaaactctcaaggatcttaccgctgttgagatccagttcgatgtaacccactcgtgcacccaactgatcttcagcatcttttactttcaccagcgtttctgggtgagcaaaaacaggaaggcaaaatgccgcaaaaaagggaataagggcgacacggaaatgttgaatactcatactcttcctttttcaatattattgaagcatttatcagggttattgtctcatgagcggatacatatttgaatgtatttagaaaaataaacaaataggggttccgcgcacatttccccgaaaagtgccacctgacgtctaagaaaccattattatcatgacattaacctataaaaataggcgtatcacgaggccctttcgtcttcaagaattaattcggtcgaaaaaagaaaaggagagggccaagagggagggcattggtgactattgagcacgtgagtatacgtgattaagcacacaaaggcagcttggagtatgtctgttattaatttcacaggtagttctggtccattggtgaaagtttgcggcttgcagagcacagaggccgcagaatgtgcactagattccgatgctgacttgctgggtattatatgtgtgcccaatagaaagagaacaattgacccggttattgcaaggaaaatttcaagtcttgtaaaagcatataaaaatagttcaggcactccgaaatacttggttggcgtgtttcgtaatcaacctaaggaggatgttttggctctggtcaatgattacggcattgatatcgtccaactgcatggagatgagtcgtggcaagaataccaagagttcctcggtttgccagttattaaaagactcgtatttccaaaagactgcaacatactactcagtgcagcttcacagaaacctcattcgtttattcccttgtttgattcagaagcaggtgggacaggtgaacttttggattggaactcgatttctgactgggttggaaggcaagagagccccgag

**Y421 Plasmid**

CEN/CEN

NLS

VP16

EL222

nat1

Ori

bla

TRP/TRP

gaattcgacatggaggcccagaataccctccttgacagtcttgacgtgcgcagctcaggggcatgatgtgactgtcgcccgtacatttagcccatacatccccatgtataatcatttgcatccatacattttgatggccgcacggcgcgaagcaaaaattacggctcctcgctgcagacctgcgagcagggaaacgctcccctcacagacgcgttgaattgtccccacgccgcgcccctgtagagaaatataaaaggttaggatttgccactgaggttcttctttcatatacttccttttaaaatcttgctaggatacagttctcacatcacatccgaacataaacaaccgcatgggccctaaaaagaagcgtaaagtcgcccccccgaccgatgtcagcctgggggacgagctccacttagacggcgaggacgtggcgatggcgcatgccgacgcgctagacgatttcgatctggacatgttgggggacggggattccccgggtccgggatttaccccccacgactccgccccctacggcgctctggatatggccgacttcgagtttgagcagatgtttaccgatgcccttggaattgacgagtacggtggggaattcggggcagacgacacacgcgttgaggtgcaaccgccggcgcagtgggtcctcgacctgatcgaggccagcccgatcgcatcggtcgtgtccgatccgcgtctcgccgacaatccgctgatcgccatcaaccaggccttcaccgacctgaccggctattccgaagaagaatgcgtcggccgcaattgccgattcctggcaggttccggcaccgagccgtggctgaccgacaagatccgccaaggcgtgcgcgagcacaagccggtgctggtcgagatcctgaactacaagaaggacggcacgccgttccgcaatgccgtgctcgttgcaccgatctacgatgacgacgacgagcttctctatttcctcggcagccaggtcgaagtcgacgacgaccagcccaacatgggcatggcgcgccgcgaacgcgccgcggaaatgctcaagacgctgtcgccgcgccagctcgaggttacgacgctggtggcatcgggcttgcgcaacaaggaagtggcggcccggctcggcctgtcggagaaaaccgtcaagatgcaccgcgggctggtgatggaaaagctcaacctgaagaccagtgccgatctggtgcgcattgccgtcgaagccggaatctaaccacttctaaataagcgaatttcttatgatttatgatttttattattaaataagttataaaaaaaataagtgtatacaaattttaaagtgactcttaggttttaaaacgaaaattcttattcttgagtaactctttcctgtaggtcaggttgctttctcaggtatagtatgaggtcgctcttattgaccacacctctaccggcagatccgctagggataacggatccacagtttattcctggcatccactaaatataatggagcccgctttttaagctggcatccagaaaaaaaaagaatcccagcaccaaaatattgttttcttcaccaaccatcagttcataggtccattctcttagcgcaactacagagaacaggggcacaaacaggcaaaaaacgggcacaacctcaatggagtgatgcaacctgcctggagtaaatgatgacacaaggcaattgacccacgcatgtatctatctcattttcttacaccttctattaccttctgctctctctgatttggaaaaagctgaaaaaaaaggttgaaaccagttccctgaaattattcccctacttgactaataagtatataaagacggtaggtattgattgtaattctgtaaatctatttcttaaacttcttaaattctacttttatagttagtcttttttttagttttaaaacaccaagaacttagtttcgaataaacacacataaacaaaatgggtaccactcttgacgacacggcttaccggtaccgcaccagtgtcccgggggacgccgaggccatcgaggcactggatgggtccttcaccaccgacaccgtcttccgcgtcaccgccaccggggacggcttcaccctgcgggaggtgccggtggacccgcccctgaccaaggtgttccccgacgacgaatcggacgacgaatcggacgacggggaggacggcgacccggactcccggacgttcgtcgcgtacggggacgacggcgacctggcgggcttcgtggtcgtctcgtactccggctggaaccgccggctgaccgtcgaggacatcgaggtcgccccggagcaccgggggcacggggtcgggcgcgcgttgatggggctcgcgacggagttcgcccgcgagcggggcgccgggcacctctggctggaggtcaccaacgtcaacgcaccggcgatccacgcgtaccggcggatggggttcaccctctgcggcctggacaccgccctgtacgacggcaccgcctcggacggcgagcaggcgctctacatgagcatgccctgcccctaattaattaaggccgctagggccctgcaggagggccgcatcatgtaattagttatgtcacgcttacattcacgccctccccccacatccgctctaaccgaaaaggaaggagttagacaacctgaagtctaggtccctatttatttttttatagttatgttagtattaagaacgttatttatatttcaaatttttcttttttttctgtacagacgcgtgtacgcatgtaacattatactgaaaaccttgcttgagaaggttttgggacgctcgaaggctttaatttgcggccaagcttggcgtaatcatggtcatagctgtttcctgtgtgaaattgttatccgctcacaattccacacaacatacgagccggaagcataaagtgtaaagcctggggtgcctaatgagtgagctaactcacattaattgcgttgcgctcactgcccgctttccagtcgggaaacctgtcgtgccagctgcattaatgaatcggccaacgcgcggggagaggcggtttgcgtattgggcgctcttccgcttcctcgctcactgactcgctgcgctcggtcgttcggctgcggcgagcggtatcagctcactcaaaggcggtaatacggttatccacagaatcaggggataacgcaggaaagaacatgtgagcaaaaggccagcaaaaggccaggaaccgtaaaaaggccgcgttgctggcgtttttccataggctccgcccccctgacgagcatcacaaaaatcgacgctcaagtcagaggtggcgaaacccgacaggactataaagataccaggcgtttccccctggaagctccctcgtgcgctctcctgttccgaccctgccgcttaccggatacctgtccgcctttctcccttcgggaagcgtggcgctttctcatagctcacgctgtaggtatctcagttcggtgtaggtcgttcgctccaagctgggctgtgtgcacgaaccccccgttcagcccgaccgctgcgccttatccggtaactatcgtcttgagtccaacccggtaagacacgacttatcgccactggcagcagccactggtaacaggattagcagagcgaggtatgtaggcggtgctacagagttcttgaagtggtggcctaactacggctacactagaaggacagtatttggtatctgcgctctgctgaagccagttaccttcggaaaaagagttggtagctcttgatccggcaaacaaaccaccgctggtagcggtggtttttttgtttgcaagcagcagattacgcgcagaaaaaaaggatctcaagaagatcctttgatcttttctacggggtctgacgctcagtggaacgaaaactcacgttaagggattttggtcatgagattatcaaaaaggatcttcacctagatccttttaaattaaaaatgaagttttaaatcaatctaaagtatatatgagtaaacttggtctgacagttaccaatgcttaatcagtgaggcacctatctcagcgatctgtctatttcgttcatccatagttgcctgactccccgtcgtgtagataactacgatacgggagggcttaccatctggccccagtgctgcaatgataccgcgagacccacgctcaccggctccagatttatcagcaataaaccagccagccggaagggccgagcgcagaagtggtcctgcaactttatccgcctccatccagtctattaattgttgccgggaagctagagtaagtagttcgccagttaatagtttgcgcaacgttgttgccattgctacaggcatcgtggtgtcacgctcgtcgtttggtatggcttcattcagctccggttcccaacgatcaaggcgagttacatgatcccccatgttgtgcaaaaaagcggttagctccttcggtcctccgatcgttgtcagaagtaagttggccgcagtgttatcactcatggttatggcagcactgcataattctcttactgtcatgccatccgtaagatgcttttctgtgactggtgagtactcaaccaagtcattctgagaatagtgtatgcggcgaccgagttgctcttgcccggcgtcaatacgggataataccgcgccacatagcagaactttaaaagtgctcatcattggaaaacgttcttcggggcgaaaactctcaaggatcttaccgctgttgagatccagttcgatgtaacccactcgtgcacccaactgatcttcagcatcttttactttcaccagcgtttctgggtgagcaaaaacaggaaggcaaaatgccgcaaaaaagggaataagggcgacacggaaatgttgaatactcatactcttcctttttcaatattattgaagcatttatcagggttattgtctcatgagcggatacatatttgaatgtatttagaaaaataaacaaataggggttccgcgcacatttccccgaaaagtgccacctgacgtctaagaaaccattattatcatgacattaacctataaaaataggcgtatcacgaggccctttcgtcttcaagaattaattcggtcgaaaaaagaaaaggagagggccaagagggagggcattggtgactattgagcacgtgagtatacgtgattaagcacacaaaggcagcttggagtatgtctgttattaatttcacaggtagttctggtccattggtgaaagtttgcggcttgcagagcacagaggccgcagaatgtgcactagattccgatgctgacttgctgggtattatatgtgtgcccaatagaaagagaacaattgacccggttattgcaaggaaaatttcaagtcttgtaaaagcatataaaaatagttcaggcactccgaaatacttggttggcgtgtttcgtaatcaacctaaggaggatgttttggctctggtcaatgattacggcattgatatcgtccaactgcatggagatgagtcgtggcaagaataccaagagttcctcggtttgccagttattaaaagactcgtatttccaaaagactgcaacatactactcagtgcagcttcacagaaacctcattcgtttattcccttgtttgattcagaagcaggtgggacaggtgaacttttggattggaactcgatttctgactgggttggaaggcaagagagccccgagagcttacattttatgttagctggtggactgacgccagaaaatgttggtgatgcgcttagattaaatggcgttattggtgttgatgtaagcggaggtgtggagacaaatggtgtaaaagactctaacaaaatagcaaatttcgtcaaaaatgctaagaaataggttattactgagtagtatttatttaagtattgtttgtgcacttgcctgcaagccttttgaaaagcaagcataaaagatctaaacataaaatctgtaaaataacaagatgtaaagataatgctaaatcatttggctttttgattgattgtacaggaaaatatacatcgcagggggttgacttttaccatttcaccgcaatggaatcaaacttgttgaagagaatgttcacaggcgcatacgctacaatgacccgattcttgctagccttttctcggtcttgcaaacaaccgccggcagcttagtatataaatacacatgtacatacctctctccgtatcctcgtaatcattttcttgtatttatcgtcttttcgctgtaaaaactttatcacacttatctcaaatacacttattaaccgcttttactattatcttctacgctgacagtaatatcaaacagtgacacatattaaacacagtggtttctttgcataaacaccatcagcctcaagtcgtcaagtaaagatttcgtgttcatgcagatagataacaatctatatgttgataattagcgttgcctcatcaatgcgagatccgtttaaccggaccctagtgcacttaccccacgttcggtccactgtgtgccgaacatgctccttcactattttaacatgtggaattaattctcatgtttgacagcttatcatcgaactctaagaggtgatacttatttactgtaaaactgtgacgataaaaccggaaggaagaataagaaaactcgaactgatctataatgcctattttctgtaaagagtttaagctatgaaagcctcggcattttggccgctcctaggtagtgctttttttccaaggacaaaacagtttctttttcttgagcaggttttatgtttcggtaatcataaacaataaataaattatttcatttatgtttaaaaataaaaaataaaaaagtattttaaatttttaaaaaagttgattataagcatgtgaccttttgcaagcaattaaattttgcaatttgtgattttaggcaaaagttacaatttctggctcgtgtaatatatgtatgctaaagtgaacttttacaaagtcgatatggacttagtcaaaagaaattttcttaaaaatatatagcactagccaatttagcacttctttatgagatatattatagactttattaagccagatttgtgtattatatgtatttacccggcgaatcatggacatacattctgaaataggtaatattctctatggtgagacagcatagataacctaggatacaagttaaaagctagtactgttttgcagtaatttttttcttttttataagaatgttaccacctaaataagttataaagtcaatagttaagtttgatatttgattgtaaaataccgtaatatatttgcatgatcaaaaggctcaatgttgactagccagcatgtcaaccactatattgatcaccgatatatggacttccacaccaactagtaatatgacaataaattcaagatattcttcatgagaatggcccagcgatatatgcggtgtgaaataccgcacagatgcgtaaggagaaaataccgcatcaggcgccattcgccattcaggctgcgcaactgttgggaagggcgatcggtgcgggcctcttcgctattacgccagctggcgaaagggggatgtgctgcaaggcgattaagttgggtaacgccagggttttcccagtcacgacgttgtaaaacgacggccagt

**PHO4^E259L^-EL222**

CEN/CEN

NLS

PHO4

E259L Mutation

Linker

EL222

nat1

Ori

bla

TRP/TRP

agcttacattttatgttagctggtggactgacgccagaaaatgttggtgatgcgcttagattaaatggcgttattggtgttgatgtaagcggaggtgtggagacaaatggtgtaaaagactctaacaaaatagcaaatttcgtcaaaaatgctaagaaataggttattactgagtagtatttatttaagtattgtttgtgcacttgcctgcaagccttttgaaaagcaagcataaaagatctaaacataaaatctgtaaaataacaagatgtaaagataatgctaaatcatttggctttttgattgattgtacaggaaaatatacatcgcagggggttgacttttaccatttcaccgcaatggaatcaaacttgttgaagagaatgttcacaggcgcatacgctacaatgacccgattcttgctagccttttctcggtcttgcaaacaaccgccggcagcttagtatataaatacacatgtacatacctctctccgtatcctcgtaatcattttcttgtatttatcgtcttttcgctgtaaaaactttatcacacttatctcaaatacacttattaaccgcttttactattatcttctacgctgacagtaatatcaaacagtgacacatattaaacacagtggtttctttgcataaacaccatcagcctcaagtcgtcaagtaaagatttcgtgttcatgcagatagataacaatctatatgttgataattagcgttgcctcatcaatgcgagatccgtttaaccggaccctagtgcacttaccccacgttcggtccactgtgtgccgaacatgctccttcactattttaacatgtggaattaattctcatgtttgacagcttatcatcgaactctaagaggtgatacttatttactgtaaaactgtgacgataaaaccggaaggaagaataagaaaactcgaactgatctataatgcctattttctgtaaagagtttaagctatgaaagcctcggcattttggccgctcctaggtagtgctttttttccaaggacaaaacagtttctttttcttgagcaggttttatgtttcggtaatcataaacaataaataaattatttcatttatgtttaaaaataaaaaataaaaaagtattttaaatttttaaaaaagttgattataagcatgtgaccttttgcaagcaattaaattttgcaatttgtgattttaggcaaaagttacaatttctggctcgtgtaatatatgtatgctaaagtgaacttttacaaagtcgatatggacttagtcaaaagaaattttcttaaaaatatatagcactagccaatttagcacttctttatgagatatattatagactttattaagccagatttgtgtattatatgtatttacccggcgaatcatggacatacattctgaaataggtaatattctctatggtgagacagcatagataacctaggatacaagttaaaagctagtactgttttgcagtaatttttttcttttttataagaatgttaccacctaaataagttataaagtcaatagttaagtttgatatttgattgtaaaataccgtaatatatttgcatgatcaaaaggctcaatgttgactagccagcatgtcaaccactatattgatcaccgatatatggacttccacaccaactagtaatatgacaataaattcaagatattcttcatgagaatggcccagcgatatatgcggtgtgaaataccgcacagatgcgtaaggagaaaataccgcatcaggcgccattcgccattcaggctgcgcaactgttgggaagggcgatcggtgcgggcctcttcgctattacgccagctggcgaaagggggatgtgctgcaaggcgattaagttgggtaacgccagggttttcccagtcacgacgttgtaaaacgacggccagtgagttcgacatggaggcccagaataccctccttgacagtcttgacgtgcgcagctcaggggcatgatgtgactgtcgcccgtacatttagcccatacatccccatgtataatcatttgcatccatacattttgatggccgcacggcgcgaagcaaaaattacggctcctcgctgcagacctgcgagcagggaaacgctcccctcacagacgcgttgaattgtccccacgccgcgcccctgtagagaaatataaaaggttaggatttgccactgaggttcttctttcatatacttccttttaaaatcttgctaggatacagttctcacatcacatccgaacataaacaaccgcatgggccctaaaaagaagcgtaaagtcgaattcggccgtacaacttctgagggaatacacggttttgtggacgatctggagcccaagagcagcattcttgataaagtcggagactttatcaccgtaaacacgaaacggcatgatgggcgcgaggacttcaacgagcaaaacgacgagctgaacagtcaagagaaccacaacagcagtgagaatgggaacgagaatgaaaatgaacaagacagtctcgcgttggacgacctagaccgcgcctttgagctggtggaaggtatggatatggactggatgatgccctcgcatgcgcaccactccccagctacaactgctacaatcaagccgcggctattatattcgccgctaatacacacgcaaagtgcggttcccgtaaccatttcgccgaacttggtcgctactgctacttccaccacatccgctaacaaagtcactaaaaacaagagtaatagtagtccgtatttgaacaagcgcagaggtaaacccgggccggattcggccacttcgctgttcgaattgcccgacagcgttatcccaactccgaaaccgaaaccgaaaccaaagcaatatccgaaagttattctgccgtcgaacagcacaagacgcgtatcaccggtcacggccaagaccagcagcagcgcagaaggcgtggtcgtagcaagtgagtctcctgtaatcgcgccgcacggatcgagccattcgcggtcgctgagtaagcgacggtcatcgggcgcgctcgtggacgatgacaagcgcgaatcacacaagcatgcactgcaagcacggcgtaatcgattagcggtcgcgctgcacgaactggcgtctttaatccccgcggagtggaaacagcaaaatgtgtcggccgcgccgtccaaagcgaccaccgtggaggcggcctgccggtacatccgtcacctacagcagaacgtgagcacgtctagaggtggaggaggctctggtggaggcggtagcggaggcggagggtcgggtggcggtggctcgggcggaggtgggtcgggtggcggcggatcaggtggaggaggctctggtggaggcggtagcggaggcggagggtcgtccggaggggcagacgacacacgcgttgaggtgcaaccgccggcgcagtgggtcctcgacctgatcgaggccagcccgatcgcatcggtcgtgtccgatccgcgtctcgccgacaatccgctgatcgccatcaaccaggccttcaccgacctgaccggctattccgaagaagaatgcgtcggccgcaattgccgattcctggcaggttccggcaccgagccgtggctgaccgacaagatccgccaaggcgtgcgcgagcacaagccggtgctggtcgagatcctgaactacaagaaggacggcacgccgttccgcaatgccgtgctcgttgcaccgatctacgatgacgacgacgagcttctctatttcctcggcagccaggtcgaagtcgacgacgaccagcccaacatgggcatggcgcgccgcgaacgcgccgcggaaatgctcaagacgctgtcgccgcgccagctcgaggttacgacgctggtggcatcgggcttgcgcaacaaggaagtggcggcccggctcggcctgtcggagaaaaccgtcaagatgcaccgcgggctggtgatggaaaagctcaacctgaagaccagtgccgatctggtgcgcattgccgtcgaagccggaatctaaccacttctaaataagcgaatttcttatgatttatgatttttattattaaataagttataaaaaaaataagtgtatacaaattttaaagtgactcttaggttttaaaacgaaaattcttattcttgagtaactctttcctgtaggtcaggttgctttctcaggtatagtatgaggtcgctcttattgaccacacctctaccggcagatccgctagggataacgcggccgcctggatccacagtttattcctggcatccactaaatataatggagcccgctttttaagctggcatccagaaaaaaaaagaatcccagcaccaaaatattgttttcttcaccaaccatcagttcataggtccattctcttagcgcaactacagagaacaggggcacaaacaggcaaaaaacgggcacaacctcaatggagtgatgcaacctgcctggagtaaatgatgacacaaggcaattgacccacgcatgtatctatctcattttcttacaccttctattaccttctgctctctctgatttggaaaaagctgaaaaaaaaggttgaaaccagttccctgaaattattcccctacttgactaataagtatataaagacggtaggtattgattgtaattctgtaaatctatttcttaaacttcttaaattctacttttatagttagtcttttttttagttttaaaacaccaagaacttagtttcgaataaacacacataaacaaaatgggtaccactcttgacgacacggcttaccggtaccgcaccagtgtcccgggggacgccgaggccatcgaggcactggatgggtccttcaccaccgacaccgtcttccgcgtcaccgccaccggggacggcttcaccctgcgggaggtgccggtggacccgcccctgaccaaggtgttccccgacgacgaatcggacgacgaatcggacgacggggaggacggcgacccggactcccggacgttcgtcgcgtacggggacgacggcgacctggcgggcttcgtggtcgtctcgtactccggctggaaccgccggctgaccgtcgaggacatcgaggtcgccccggagcaccgggggcacggggtcgggcgcgcgttgatggggctcgcgacggagttcgcccgcgagcggggcgccgggcacctctggctggaggtcaccaacgtcaacgcaccggcgatccacgcgtaccggcggatggggttcaccctctgcggcctggacaccgccctgtacgacggcaccgcctcggacggcgagcaggcgctctacatgagcatgccctgcccctaattaattaaggccgctagggccctgcaggagggccgcatcatgtaattagttatgtcacgcttacattcacgccctccccccacatccgctctaaccgaaaaggaaggagttagacaacctgaagtctaggtccctatttatttttttatagttatgttagtattaagaacgttatttatatttcaaatttttcttttttttctgtacagacgcgtgtacgcatgtaacattatactgaaaaccttgcttgagaaggttttgggacgctcgaaggctttaatttgcggccaagcttggcgtaatcatggtcatagctgtttcctgtgtgaaattgttatccgctcacaattccacacaacatacgagccggaagcataaagtgtaaagcctggggtgcctaatgagtgagctaactcacattaattgcgttgcgctcactgcccgctttccagtcgggaaacctgtcgtgccagctgcattaatgaatcggccaacgcgcggggagaggcggtttgcgtattgggcgctcttccgcttcctcgctcactgactcgctgcgctcggtcgttcggctgcggcgagcggtatcagctcactcaaaggcggtaatacggttatccacagaatcaggggataacgcaggaaagaacatgtgagcaaaaggccagcaaaaggccaggaaccgtaaaaaggccgcgttgctggcgtttttccataggctccgcccccctgacgagcatcacaaaaatcgacgctcaagtcagaggtggcgaaacccgacaggactataaagataccaggcgtttccccctggaagctccctcgtgcgctctcctgttccgaccctgccgcttaccggatacctgtccgcctttctcccttcgggaagcgtggcgctttctcatagctcacgctgtaggtatctcagttcggtgtaggtcgttcgctccaagctgggctgtgtgcacgaaccccccgttcagcccgaccgctgcgccttatccggtaactatcgtcttgagtccaacccggtaagacacgacttatcgccactggcagcagccactggtaacaggattagcagagcgaggtatgtaggcggtgctacagagttcttgaagtggtggcctaactacggctacactagaaggacagtatttggtatctgcgctctgctgaagccagttaccttcggaaaaagagttggtagctcttgatccggcaaacaaaccaccgctggtagcggtggtttttttgtttgcaagcagcagattacgcgcagaaaaaaaggatctcaagaagatcctttgatcttttctacggggtctgacgctcagtggaacgaaaactcacgttaagggattttggtcatgagattatcaaaaaggatcttcacctagatccttttaaattaaaaatgaagttttaaatcaatctaaagtatatatgagtaaacttggtctgacagttaccaatgcttaatcagtgaggcacctatctcagcgatctgtctatttcgttcatccatagttgcctgactccccgtcgtgtagataactacgatacgggagggcttaccatctggccccagtgctgcaatgataccgcgagacccacgctcaccggctccagatttatcagcaataaaccagccagccggaagggccgagcgcagaagtggtcctgcaactttatccgcctccatccagtctattaattgttgccgggaagctagagtaagtagttcgccagttaatagtttgcgcaacgttgttgccattgctacaggcatcgtggtgtcacgctcgtcgtttggtatggcttcattcagctccggttcccaacgatcaaggcgagttacatgatcccccatgttgtgcaaaaaagcggttagctccttcggtcctccgatcgttgtcagaagtaagttggccgcagtgttatcactcatggttatggcagcactgcataattctcttactgtcatgccatccgtaagatgcttttctgtgactggtgagtactcaaccaagtcattctgagaatagtgtatgcggcgaccgagttgctcttgcccggcgtcaatacgggataataccgcgccacatagcagaactttaaaagtgctcatcattggaaaacgttcttcggggcgaaaactctcaaggatcttaccgctgttgagatccagttcgatgtaacccactcgtgcacccaactgatcttcagcatcttttactttcaccagcgtttctgggtgagcaaaaacaggaaggcaaaatgccgcaaaaaagggaataagggcgacacggaaatgttgaatactcatactcttcctttttcaatattattgaagcatttatcagggttattgtctcatgagcggatacatatttgaatgtatttagaaaaataaacaaataggggttccgcgcacatttccccgaaaagtgccacctgacgtctaagaaaccattattatcatgacattaacctataaaaataggcgtatcacgaggccctttcgtcttcaagaattaattcggtcgaaaaaagaaaaggagagggccaagagggagggcattggtgactattgagcacgtgagtatacgtgattaagcacacaaaggcagcttggagtatgtctgttattaatttcacaggtagttctggtccattggtgaaagtttgcggcttgcagagcacagaggccgcagaatgtgcactagattccgatgctgacttgctgggtattatatgtgtgcccaatagaaagagaacaattgacccggttattgcaaggaaaatttcaagtcttgtaaaagcatataaaaatagttcaggcactccgaaatacttggttggcgtgtttcgtaatcaacctaaggaggatgttttggctctggtcaatgattacggcattgatatcgtccaactgcatggagatgagtcgtggcaagaataccaagagttcctcggtttgccagttattaaaagactcgtatttccaaaagactgcaacatactactcagtgcagcttcacagaaacctcattcgtttattcccttgtttgattcagaagcaggtgggacaggtgaacttttggattggaactcgatttctgactgggttggaaggcaagagagccccgag

**UME6^C770A/C787A^-EL222**

CEN/CEN

NLS

UME6

C770A/C787A Mutation

Linker

EL222

nat1

Ori

bla

TRP/TRP

agcttacattttatgttagctggtggactgacgccagaaaatgttggtgatgcgcttagattaaatggcgttattggtgttgatgtaagcggaggtgtggagacaaatggtgtaaaagactctaacaaaatagcaaatttcgtcaaaaatgctaagaaataggttattactgagtagtatttatttaagtattgtttgtgcacttgcctgcaagccttttgaaaagcaagcataaaagatctaaacataaaatctgtaaaataacaagatgtaaagataatgctaaatcatttggctttttgattgattgtacaggaaaatatacatcgcagggggttgacttttaccatttcaccgcaatggaatcaaacttgttgaagagaatgttcacaggcgcatacgctacaatgacccgattcttgctagccttttctcggtcttgcaaacaaccgccggcagcttagtatataaatacacatgtacatacctctctccgtatcctcgtaatcattttcttgtatttatcgtcttttcgctgtaaaaactttatcacacttatctcaaatacacttattaaccgcttttactattatcttctacgctgacagtaatatcaaacagtgacacatattaaacacagtggtttctttgcataaacaccatcagcctcaagtcgtcaagtaaagatttcgtgttcatgcagatagataacaatctatatgttgataattagcgttgcctcatcaatgcgagatccgtttaaccggaccctagtgcacttaccccacgttcggtccactgtgtgccgaacatgctccttcactattttaacatgtggaattaattctcatgtttgacagcttatcatcgaactctaagaggtgatacttatttactgtaaaactgtgacgataaaaccggaaggaagaataagaaaactcgaactgatctataatgcctattttctgtaaagagtttaagctatgaaagcctcggcattttggccgctcctaggtagtgctttttttccaaggacaaaacagtttctttttcttgagcaggttttatgtttcggtaatcataaacaataaataaattatttcatttatgtttaaaaataaaaaataaaaaagtattttaaatttttaaaaaagttgattataagcatgtgaccttttgcaagcaattaaattttgcaatttgtgattttaggcaaaagttacaatttctggctcgtgtaatatatgtatgctaaagtgaacttttacaaagtcgatatggacttagtcaaaagaaattttcttaaaaatatatagcactagccaatttagcacttctttatgagatatattatagactttattaagccagatttgtgtattatatgtatttacccggcgaatcatggacatacattctgaaataggtaatattctctatggtgagacagcatagataacctaggatacaagttaaaagctagtactgttttgcagtaatttttttcttttttataagaatgttaccacctaaataagttataaagtcaatagttaagtttgatatttgattgtaaaataccgtaatatatttgcatgatcaaaaggctcaatgttgactagccagcatgtcaaccactatattgatcaccgatatatggacttccacaccaactagtaatatgacaataaattcaagatattcttcatgagaatggcccagcgatatatgcggtgtgaaataccgcacagatgcgtaaggagaaaataccgcatcaggcgccattcgccattcaggctgcgcaactgttgggaagggcgatcggtgcgggcctcttcgctattacgccagctggcgaaagggggatgtgctgcaaggcgattaagttgggtaacgccagggttttcccagtcacgacgttgtaaaacgacggccagtgagttcgacatggaggcccagaataccctccttgacagtcttgacgtgcgcagctcaggggcatgatgtgactgtcgcccgtacatttagcccatacatccccatgtataatcatttgcatccatacattttgatggccgcacggcgcgaagcaaaaattacggctcctcgctgcagacctgcgagcagggaaacgctcccctcacagacgcgttgaattgtccccacgccgcgcccctgtagagaaatataaaaggttaggatttgccactgaggttcttctttcatatacttccttttaaaatcttgctaggatacagttctcacatcacatccgaacataaacaaccgcatgggccctaaaaagaagcgtaaagtcgaattgctagacaaggcgcgctctcaaagcaaacacatggacgaatctaatgcggctgcctctctgctttcgatggaaacaaccgccaacaatcatcactatttgcacaataaaacatctcgtgccacgctgatgaatagcagccaagacggcaaaaaacatgcagaagatgaagttagtgatggagctaactcccgccatcctacaatttccagtgctagcatcgaatctctcaagacaacctacgatgaaaaccctttgctttctattatgaaatcgacatgtgcgcccaacaacactcccgtgcatactccgtctggttcgccgagtttgaaagtccaaagtggcggagatatcaaagacgatcctaaggaaaacgatactactactacgaccaatactacacttcaagatcgccgtgacagcgataacgctgtgcatgctgccgccagcccactcgcgccttccaatacaccttcggatcctaagtcattgtgcaatggccatgtcgcacaggctacagacccacaaatttccggtgctattcagccgcagtatactgcaaccaacgaggatgttttcccttactcctccacctccactaatagtaacactgccactactactatcgtcgccggcgccaaaaaaaaaatacatttgccgccaccacaagctccagcggtttcttcccccggtaccaccgcagcaggctcgggcgcgggcacgggctcgggcatccgttcccgcacaggatcggatttgccgctcatcattaccagcgccaacaagaacaacggtaagactaccaattcgcctatgtcgatactgagcagaaacaacagtaccaacaacaacgacaataattcgatacaaagctcagattcgagagaatcctctaataacaatgagattggcgggtatttgcgcggcggaactaagcgcggcggcagtccatctaacgactctcaggtccagcataatgtgcatgatgaccaatgtgccgtgggcgtggcgcccaggaacttctatttcaacaaggatagagagataacagacccaaatgtaaaactggacgagaacgaatcaaaaatcaacatatcgttctggctaaattcgaaatacagagatgaggcttattctttgaatgaatcatcctccaacaatgctagttcaaacacggatacgcctacaaactctcgacatgcgaacaccagctcttccattaccagcagaaacaatttccagcattttaggttcaaccaaataccttctcaacctccaacttccgcatcttcgtttacaagcacaaacaacaacaaccctcaacggaacaatatcaatcgcggtgaagacccgtttgccacttcgtcaagaccttctactgggtttttttacggcgatttgccgaatcgtaacaatagaaatagtcccttccatacaaatgaacaatacatcccaccacctccaccgaaatacatcaattctaaattggatggattgagatcaagattattgctcggtccgaattctgcatcttcatctaccaaactagacgacgacttgggtacagcagcagcagtgctatcaaacatgagatcatccccatatagaactcatgataaacccatttccaatgtcaatgacatgaataacacaaatgcgctcggtgtgccggctagtaggcctcattcgtcatcttttccatcaaagggtgtcttaagaccaattctgttacgtatccataattccgaacaacaacccattttcgaaagcaacaattctacagctgtttttgatgaagaccaggaccaaaatcaagacttgtctccttaccatttaaatctaaactctaaaaaggttttagatcccacttttgagtcaaggacaaggcaagttacttggaataagaatggtaagcgaatagacagacgcctttctgctccagaacaacaacagcaactggaagttccaccattgaaaaaatcgagaaggtcagttggaaacgcaagagtagcaagccaaaccaatagcgattataattctcttggcgaatcctcaacttcgtcagctccatcgtctccatctttgaaggcttcttctggcttggcatataccgctgattatcctaacgctacttcgccggatttcgctaaatctaaaggaaaaaatgtcaagcctaaggcaaaatcaaaggcgaaacagtcatcaaagaaaagaccaaataatactacttcgaaatcaaaagcaaacaattctcaagaatcgaataatgctacttcctcaacgtctcaaggtacaaggtcccgtactggtgcctggatttgtagattaaggaaaaagaagtgtaccgaggaaagaccgcacgctttcaactgtgaaaggttgaaattggactgtcactatgacgcgttcaaaccagattttgtatctgatccaaagaaaaaacagatgaaactggaggaaatcaagaaaaaaacaaaagaggccaaaagaagagcaatgaaaaaaaaatctagaggtggaggaggctctggtggaggcggtagcggaggcggagggtcgggtggcggtggctcgggcggaggtgggtcgggtggcggcggatcaggtggaggaggctctggtggaggcggtagcggaggcggagggtcgtccggaggggcagacgacacacgcgttgaggtgcaaccgccggcgcagtgggtcctcgacctgatcgaggccagcccgatcgcatcggtcgtgtccgatccgcgtctcgccgacaatccgctgatcgccatcaaccaggccttcaccgacctgaccggctattccgaagaagaatgcgtcggccgcaattgccgattcctggcaggttccggcaccgagccgtggctgaccgacaagatccgccaaggcgtgcgcgagcacaagccggtgctggtcgagatcctgaactacaagaaggacggcacgccgttccgcaatgccgtgctcgttgcaccgatctacgatgacgacgacgagcttctctatttcctcggcagccaggtcgaagtcgacgacgaccagcccaacatgggcatggcgcgccgcgaacgcgccgcggaaatgctcaagacgctgtcgccgcgccagctcgaggttacgacgctggtggcatcgggcttgcgcaacaaggaagtggcggcccggctcggcctgtcggagaaaaccgtcaagatgcaccgcgggctggtgatggaaaagctcaacctgaagaccagtgccgatctggtgcgcattgccgtcgaagccggaatctaaccacttctaaataagcgaatttcttatgatttatgatttttattattaaataagttataaaaaaaataagtgtatacaaattttaaagtgactcttaggttttaaaacgaaaattcttattcttgagtaactctttcctgtaggtcaggttgctttctcaggtatagtatgaggtcgctcttattgaccacacctctaccggcagatccgctagggataacgcggccgcctggatccacagtttattcctggcatccactaaatataatggagcccgctttttaagctggcatccagaaaaaaaaagaatcccagcaccaaaatattgttttcttcaccaaccatcagttcataggtccattctcttagcgcaactacagagaacaggggcacaaacaggcaaaaaacgggcacaacctcaatggagtgatgcaacctgcctggagtaaatgatgacacaaggcaattgacccacgcatgtatctatctcattttcttacaccttctattaccttctgctctctctgatttggaaaaagctgaaaaaaaaggttgaaaccagttccctgaaattattcccctacttgactaataagtatataaagacggtaggtattgattgtaattctgtaaatctatttcttaaacttcttaaattctacttttatagttagtcttttttttagttttaaaacaccaagaacttagtttcgaataaacacacataaacaaaatgggtaccactcttgacgacacggcttaccggtaccgcaccagtgtcccgggggacgccgaggccatcgaggcactggatgggtccttcaccaccgacaccgtcttccgcgtcaccgccaccggggacggcttcaccctgcgggaggtgccggtggacccgcccctgaccaaggtgttccccgacgacgaatcggacgacgaatcggacgacggggaggacggcgacccggactcccggacgttcgtcgcgtacggggacgacggcgacctggcgggcttcgtggtcgtctcgtactccggctggaaccgccggctgaccgtcgaggacatcgaggtcgccccggagcaccgggggcacggggtcgggcgcgcgttgatggggctcgcgacggagttcgcccgcgagcggggcgccgggcacctctggctggaggtcaccaacgtcaacgcaccggcgatccacgcgtaccggcggatggggttcaccctctgcggcctggacaccgccctgtacgacggcaccgcctcggacggcgagcaggcgctctacatgagcatgccctgcccctaattaattaaggccgctagggccctgcaggagggccgcatcatgtaattagttatgtcacgcttacattcacgccctccccccacatccgctctaaccgaaaaggaaggagttagacaacctgaagtctaggtccctatttatttttttatagttatgttagtattaagaacgttatttatatttcaaatttttcttttttttctgtacagacgcgtgtacgcatgtaacattatactgaaaaccttgcttgagaaggttttgggacgctcgaaggctttaatttgcggccaagcttggcgtaatcatggtcatagctgtttcctgtgtgaaattgttatccgctcacaattccacacaacatacgagccggaagcataaagtgtaaagcctggggtgcctaatgagtgagctaactcacattaattgcgttgcgctcactgcccgctttccagtcgggaaacctgtcgtgccagctgcattaatgaatcggccaacgcgcggggagaggcggtttgcgtattgggcgctcttccgcttcctcgctcactgactcgctgcgctcggtcgttcggctgcggcgagcggtatcagctcactcaaaggcggtaatacggttatccacagaatcaggggataacgcaggaaagaacatgtgagcaaaaggccagcaaaaggccaggaaccgtaaaaaggccgcgttgctggcgtttttccataggctccgcccccctgacgagcatcacaaaaatcgacgctcaagtcagaggtggcgaaacccgacaggactataaagataccaggcgtttccccctggaagctccctcgtgcgctctcctgttccgaccctgccgcttaccggatacctgtccgcctttctcccttcgggaagcgtggcgctttctcatagctcacgctgtaggtatctcagttcggtgtaggtcgttcgctccaagctgggctgtgtgcacgaaccccccgttcagcccgaccgctgcgccttatccggtaactatcgtcttgagtccaacccggtaagacacgacttatcgccactggcagcagccactggtaacaggattagcagagcgaggtatgtaggcggtgctacagagttcttgaagtggtggcctaactacggctacactagaaggacagtatttggtatctgcgctctgctgaagccagttaccttcggaaaaagagttggtagctcttgatccggcaaacaaaccaccgctggtagcggtggtttttttgtttgcaagcagcagattacgcgcagaaaaaaaggatctcaagaagatcctttgatcttttctacggggtctgacgctcagtggaacgaaaactcacgttaagggattttggtcatgagattatcaaaaaggatcttcacctagatccttttaaattaaaaatgaagttttaaatcaatctaaagtatatatgagtaaacttggtctgacagttaccaatgcttaatcagtgaggcacctatctcagcgatctgtctatttcgttcatccatagttgcctgactccccgtcgtgtagataactacgatacgggagggcttaccatctggccccagtgctgcaatgataccgcgagacccacgctcaccggctccagatttatcagcaataaaccagccagccggaagggccgagcgcagaagtggtcctgcaactttatccgcctccatccagtctattaattgttgccgggaagctagagtaagtagttcgccagttaatagtttgcgcaacgttgttgccattgctacaggcatcgtggtgtcacgctcgtcgtttggtatggcttcattcagctccggttcccaacgatcaaggcgagttacatgatcccccatgttgtgcaaaaaagcggttagctccttcggtcctccgatcgttgtcagaagtaagttggccgcagtgttatcactcatggttatggcagcactgcataattctcttactgtcatgccatccgtaagatgcttttctgtgactggtgagtactcaaccaagtcattctgagaatagtgtatgcggcgaccgagttgctcttgcccggcgtcaatacgggataataccgcgccacatagcagaactttaaaagtgctcatcattggaaaacgttcttcggggcgaaaactctcaaggatcttaccgctgttgagatccagttcgatgtaacccactcgtgcacccaactgatcttcagcatcttttactttcaccagcgtttctgggtgagcaaaaacaggaaggcaaaatgccgcaaaaaagggaataagggcgacacggaaatgttgaatactcatactcttcctttttcaatattattgaagcatttatcagggttattgtctcatgagcggatacatatttgaatgtatttagaaaaataaacaaataggggttccgcgcacatttccccgaaaagtgccacctgacgtctaagaaaccattattatcatgacattaacctataaaaataggcgtatcacgaggccctttcgtcttcaagaattaattcggtcgaaaaaagaaaaggagagggccaagagggagggcattggtgactattgagcacgtgagtatacgtgattaagcacacaaaggcagcttggagtatgtctgttattaatttcacaggtagttctggtccattggtgaaagtttgcggcttgcagagcacagaggccgcagaatgtgcactagattccgatgctgacttgctgggtattatatgtgtgcccaatagaaagagaacaattgacccggttattgcaaggaaaatttcaagtcttgtaaaagcatataaaaatagttcaggcactccgaaatacttggttggcgtgtttcgtaatcaacctaaggaggatgttttggctctggtcaatgattacggcattgatatcgtccaactgcatggagatgagtcgtggcaagaataccaagagttcctcggtttgccagttattaaaagactcgtatttccaaaagactgcaacatactactcagtgcagcttcacagaaacctcattcgtttattcccttgtttgattcagaagcaggtgggacaggtgaacttttggattggaactcgatttctgactgggttggaaggcaagagagccccgag
